# Cryo-EM reconstructions of inhibitor-bound SMG1 kinase reveal an autoinhibitory state dependent on SMG8

**DOI:** 10.1101/2021.07.28.454180

**Authors:** Lukas M. Langer, Fabien Bonneau, Yair Gat, Elena Conti

## Abstract

The PI3K-related kinase (PIKK) SMG1 monitors progression of metazoan nonsense-mediated mRNA decay (NMD) by phosphorylating the RNA helicase UPF1. Previous work has shown that the activity of SMG1 is impaired by small molecule inhibitors, is reduced by the SMG1 interactors SMG8 and SMG9, and is downregulated by the so-called SMG1 insertion domain. However, the molecular basis for this complex regulatory network has remained elusive. Here, we present cryo-electron microscopy reconstructions of human SMG1-9 and SMG1-8-9 complexes bound to either a SMG1 inhibitor or a non-hydrolyzable ATP analogue at overall resolutions ranging from 2.8 to 3.6 Å. These structures reveal the basis with which a small molecule inhibitor preferentially targets SMG1 over other PIKKs. By comparison with our previously reported substrate-bound structure (Langer et al. 2020), we show that the SMG1 insertion domain can exert an autoinhibitory function by directly blocking the substrate binding path as well as overall access to the SMG1 kinase active site. Together with biochemical analysis, our data indicate that SMG1 autoinhibition is stabilized by the presence of SMG8. Our results explain the specific inhibition of SMG1 by an ATP-competitive small molecule, provide insights into regulation of its kinase activity within the NMD pathway, and expand the understanding of PIKK regulatory mechanisms in general.

## Introduction

Nonsense-mediated mRNA decay (NMD) is a co-translational mRNA quality control pathway central to the detection and removal of mRNAs containing premature termination codons as well as to the regulation of many physiological transcripts (Kurosaki, Popp, and Maquat 2019; Karousis and Muhlemann 2019). In metazoans, translation termination at a premature stop codon triggers phosphorylation of the RNA helicase UPF1, which then enables the recruitment of downstream effectors, in turn resulting in the degradation of the targeted mRNA by ribonucleases (Ohnishi et al. 2003; Yamashita 2013; Chakrabarti et al. 2014; Nicholson et al. 2014; Okada-Katsuhata et al. 2012; Kashima et al. 2006). UPF1 phosphorylation is thus a crucial point in metazoan NMD; it occurs specifically at particular Ser-containing motifs and is mediated by the SMG1 kinase (Denning et al. 2001; Yamashita et al. 2001).

SMG1 is a member of the phosphatidylinositol-3-kinase-related kinase (PIKK) family, which also includes the ATM, ATR, DNA-PKc and mTOR kinases (Keith and Schreiber 1995). Despite the confounding name reflecting a possible evolutionary origin with a lipid kinase (Keith and Schreiber 1995), and in retrospect the ability of some PIKK family members to bind inositol-6-phosphate (Gat et al. 2019), all active PIKKs are Ser-Thr protein kinases. These large, multidomain enzymes share an overall similar architecture: a C-terminal catalytic module (comprising the so-called FAT, kinase and FATC domains) and an N-terminal α-solenoid (that typically serves as a protein-protein interaction module). Indeed, PIKKs bind to and are regulated by interacting proteins, with which they form large and dynamic assemblies. In addition, some PIKKs contain an autoregulatory element, particularly in the variable region connecting the kinase and FATC domains, that is known as the PIKK-regulatory domain (PRD) (Baretic and Williams 2014; Imseng, Aylett, and Maier 2018; Jansma and Hopfner 2020). Finally, the kinase domains of PIKKs have been the target of numerous efforts in the development of specific inhibitors. Not surprisingly, given the central roles of these kinases in pathways surveilling cellular homeostasis, small molecules targeting specific PIKKs have been approved as therapeutics or are being evaluated in clinical trials (Janku, Yap, and Meric-Bernstam 2018; Zhang, Duan, and Zheng 2011; Durant et al. 2018). Nevertheless, the high conservation of the PIKK kinase domain has complicated efforts to develop inhibitors specifically targeting only a selected member of this family of enzymes. Structural data rationalizing binding specificity of such compounds to PIKKs are still sparse in general and entirely missing for the SMG1 kinase.

In the case of SMG1, the two interacting factors SMG8 and SMG9 have been linked to dysregulated NMD and neurodevelopmental disorders in humans (Alzahrani et al. 2020; Shaheen et al. 2016; Yamashita et al. 2009). Previous cryo-electron microscopy (cryo-EM) studies have revealed the molecular interactions that underpin the structure of the human SMG1-SMG8-SMG9 complex (Gat et al. 2019; Zhu et al. 2019) and the determinants with which it recognizes and phosphorylates UPF1 peptides with specific Leu-Ser-Gln (LSQ) motifs (Langer et al. 2020). SMG8 and SMG9 form an unusual G-domain heterodimer (Gat et al. 2019; Langer et al. 2020; Li et al. 2017). The G-domain of SMG9 binds both the *α*-solenoid and the catalytic module of SMG1 while the G-domain of SMG8 engages only the *α*-solenoid, thus rationalizing biochemical studies pointing to the crucial role of SMG9 in enabling the incorporation of SMG8 into the complex (Yamashita et al. 2009; Arias-Palomo et al. 2011). In turn, SMG8 appears to have a direct regulatory function on SMG1, as its removal or C-terminal truncation results in hyper-activation of SMG1 kinase activity (Yamashita et al. 2009; Deniaud et al. 2015; Arias-Palomo et al. 2011; Zhu et al. 2019). However, the C-terminal domain of SMG8 remains poorly defined in all available cryo-EM reconstructions (Zhu et al. 2019; Gat et al. 2019; Langer et al. 2020), hindering a molecular understanding of its regulatory role.

Another portion of the complex reported to downregulate SMG1 kinase activity is the so-called insertion domain, a large 1200-residue region connecting the SMG1 kinase and FATC domains. Removal of the SMG1 insertion domain causes hyper-activation of the kinase (Deniaud et al. 2015; Zhu et al. 2019), similarly to the effect reported for the PRDs of other PIKKs (McMahon et al. 2002; Edinger and Thompson 2004; Xiao et al. 2019; Mordes et al. 2008). However, the SMG1 insertion domain shows no sequence similarity to the PRDs of other PIKKs and is remarkably larger in comparison. None of the current cryo-EM reconstructions of SMG1 or its complexes show density corresponding to the SMG1 insertion domain (Zhu et al. 2019; Gat et al. 2019; Langer et al. 2020), and it is thus unclear how it regulates kinase activity.

Here, we used cryo-EM and biochemical approaches to study the basis with which SMG1 is specifically inhibited by a small molecule compound (SMG1**i**), and in doing so we also identified the molecular mechanisms with which SMG1 is down-regulated by its own insertion domain in *cis* and SMG8 in *trans*.

## Results

### The compound SMG1i specifically inhibits SMG1 *in vitro* by competing with ATP binding

We set out to study the inhibitory mechanism of SMG1**i** (Fig. 1 A), a small-molecule compound based on a pyrimidine derivative that has been reported to inhibit SMG1 catalytic activity (Gopalsamy et al. 2012; Mino et al. 2019). We expressed and purified human wild-type SMG1-8-9 complex from piggyBac transposase generated HEK293T stable cell pools, essentially as described before (Gat et al. 2019). We have previously shown that the use of a UPF1 peptide comprising the UPF1 phosphorylation site 1078 (UPF1-LSQ) allows monitoring of its specific phosphorylation by SMG1 over time using mass spectrometry (Langer et al. 2020). We made use of this method to assay the effect and potency of SMG1**i** (Fig. 1 and Fig. 1 Supp. 1). To avoid artefacts caused by ATP depletion, we performed an ATP titration and analyzed the end-point measurements (Fig. 1 Supp. 1 B). As a control, we used a UPF1-LDQ peptide in which we exchanged the phosphor-acceptor Ser1078 of UPF1-LSQ to an aspartic acid. Next, we repeated end-point measurements under conditions of stable ATP concentration and increasing amounts of SMG1**i** (Fig. 1 B). In this assay, we observed a significant reduction of UPF1-LSQ phosphorylation when adding SMG1**i** at concentrations equimolar to the enzyme (Fig. 1 B and Fig. 1 Supp. 1 C). To corroborate these results, we repeated titration of SMG1**i** under conditions of stable ATP concentration using a radioactivity-based phosphorylation assay and full-length UPF1 as a substrate. Again, we observed a reduction in UPF1 phosphorylation in the presence of low micromolar amounts of SMG1**i** (Fig. 1 C).

**Figure 1:**
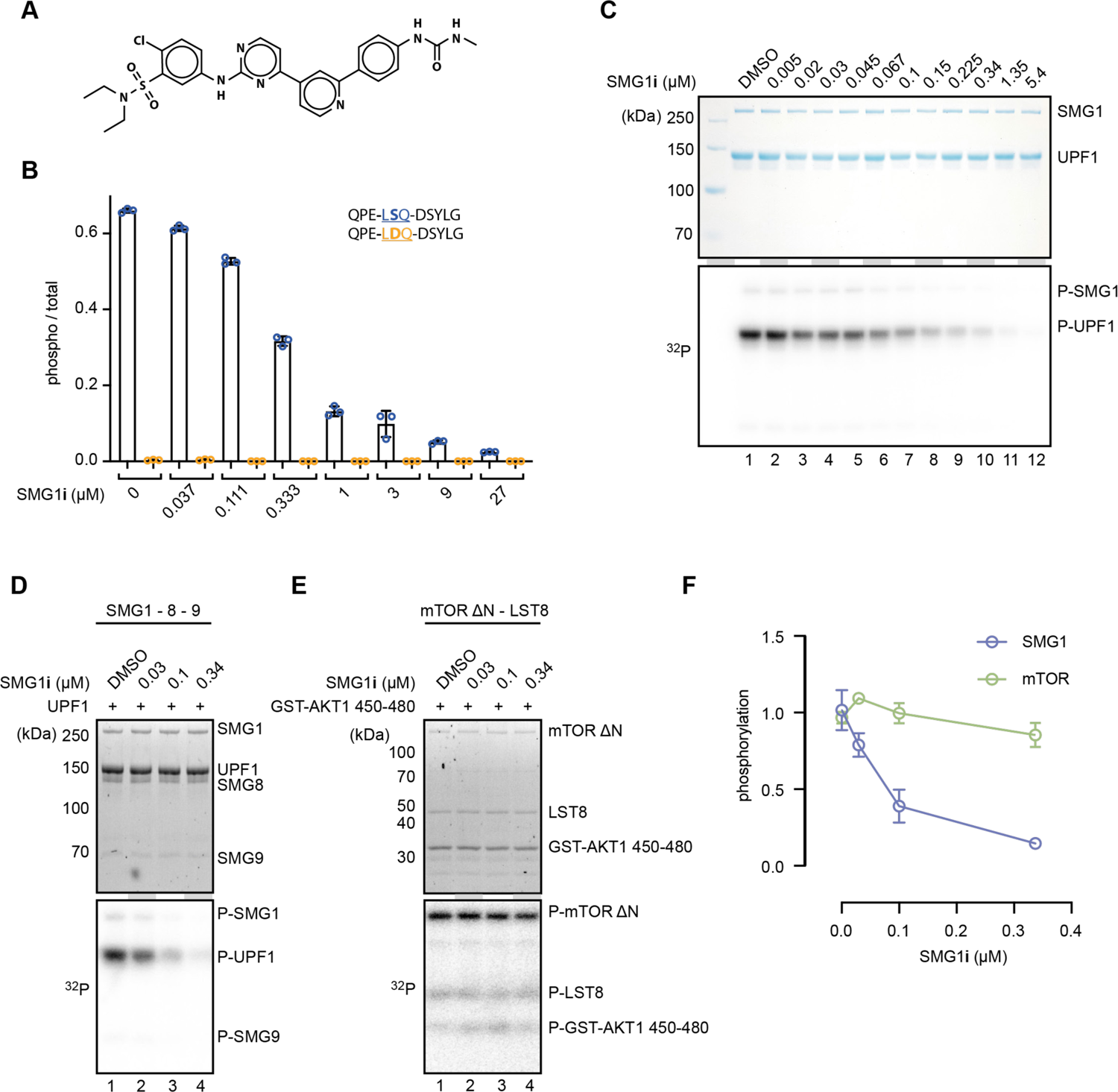
SMG1i specifically inhibits SMG1 kinase activity *in vitro*. A) Structure of the SMG1 inhibitor (SMG1**i**). B) Titration of SMG1**i** using a mass spectrometry-based phosphorylation assay with 500 nM SMG1-8-9 and the indicated UPF1-derived peptides as substrates. C) Titration of SMG1**i** using a radioactivity-based phosphorylation assay with 100 nM SMG1-8-9 and full-length UPF1 as a substrate. The Coomassie-stained gel is shown on top, and the corresponding radioactive signal below. D) and E) Titration of SMG1**i** using a radioactivity-based phosphorylation assay with either SMG1-8-9 and full-length UPF1 (D) or mTOR^ΔN^-LST8 and GST-AKT1 (E) as a substrate. Stain-free gels are shown on top and the corresponding radioactive signal at the bottom (Ladner et al. 2004). F) Quantification of normalized UPF1 or mTOR (auto-)phosphorylation in the presence of increasing amounts of SMG1**i** for both SMG1 and mTOR. For each datapoint, the mean is shown with standard deviations of the three replicates indicated.

To assess the specificity of SMG1**i**, we tested its effect on the mTOR kinase using a similar radioactive kinase assay. In this experiment, we used a truncated mTOR^ΔN^-LST8 complex previously shown to be constitutively active and responsive to mTOR inhibitors (Yang et al. 2013). As an mTOR kinase substrate, we used a GST-AKT1 ^450-480^ fusion protein. We observed that SMG1**i** only affected mTOR activity in the highest concentration range, well beyond concentrations that had an observable effect on SMG1 activity (Fig. 1 Supp. 2 A). We selected four different concentrations of SMG1**i** based on these assays (Fig. 1 C, Fig. 1 Supp. 2 A) and repeated the radioactivity-based kinase assay for both SMG1 and mTOR in triplicates (Fig. 1 D and E, Fig. 1 Supp. 2 B and C). Upon quantifying the levels of phosphorylation using densitometry and normalizing them to the amount of protein loaded in each lane, we found that SMG1**i** robustly inhibited SMG1 while mTOR activity was only weakly affected (Fig. 1 F). We concluded that SMG1**i** displays high potency and specificity in inhibiting the SMG1 kinase *in vitro*.

### Cryo-EM structure of SMG1-8-9 bound to the SMG1i inhibitor

To understand the molecular basis for the SMG1**i** mode of action, we reconstituted purified wild-type SMG1-8-9 with an excess of SMG1**i** and subjected the sample to single-particle cryo-EM analysis. The resulting reconstruction reached an overall resolution of 3.0 Å, and we observed clear density for the SMG1 inhibitor bound to the kinase active site (Fig. 2 A, Fig. 2 Supp. 1, 2 and 4 B and C). Superposition with our previously published AMPPNP- and substrate-bound model (Langer et al. 2020) revealed that SMG1**i** exploits essentially the same binding site as the ATP analogue. However, SMG1**i** wedges deeper in between the N- and C-lobe of the kinase domain and is engaged in more extensive interactions (Fig. 2 B and Fig. 2 Supp. 4 A and D), rationalizing its potent ability to compete with ATP.

**Figure 2:**
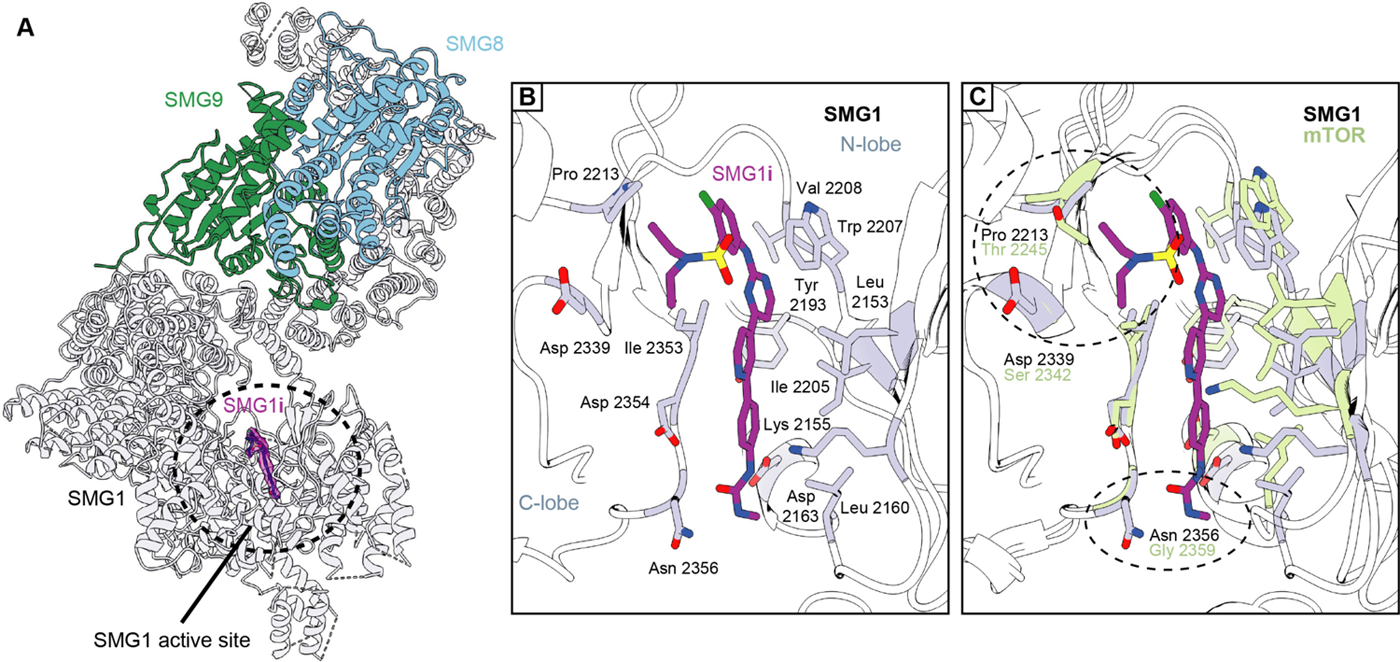
Structural basis for selective targeting of SMG1 by the SMG1 inhibitor. A) Model of the SMG1-8-9 kinase complex bound to SMG1**i**. SMG1 is in grey, SMG8 is in blue and SMG9 is shown in green. SMG1**i** is shown as a magenta model overlaid with the isolated transparent density. Approximate location of the SMG1 active site is indicated by a black circle. B) Key interactions of SMG1**i** with SMG1 active site residues. Important residues located in either the N- or the C-lobe of the SMG1 kinase domain are colored grey. Other parts of SMG1 are transparent and interactions of SMG1**i** with SMG1 backbone are not shown. C) Superposition of SMG1**i**-bound SMG1 with the mTOR active site (PDB: 4jsp) over the catalytic loop. Key SMG1 residues indicated in B) are shown alongside the respective mTOR residues colored in green. Regions possibly accounting for preferential interaction of SMG1**i** with SMG1 over mTOR are circled and the relevant residues are labeled.

While many of the SMG1 residues interacting with SMG1**i** are conserved within the PIKK family, two contact sites appear to be specific: SMG1 Pro2213 and Asp2339 contact the sulfonamide moiety of the inhibitor while Asn2356 forms hydrogen bonds with the phenyl urea/carbamide group (Fig. 2 B). Superposition of the SMG1 active site in the SMG1**i**-bound structure with the active site of mTOR^ΔN^ shows that the corresponding residues have diverged and would not be able to mediate analogous interactions with the SMG1**i** sulfonamide group (mTOR Thr 2245 and Ser2342 at the positions of SMG1 Pro2213 and Asp2339) or with the urea group (mTOR Gly2359 at the position of SMG1 Asn2356) (Fig. 2 C). The same regions of the small molecule are diverging or missing in the mTOR inhibitor Torin 2 (Fig. 2 Supp. 4 E). Together, these observations rationalize the specificity of SMG1**i** for SMG1 over mTOR^ΔN^ that we observed in the *in vitro* assays (Fig. 1 D to F).

Superposition with other PIKKs suggests the presence of potentially similar discriminatory interactions: like mTOR, DNA-PK lacks a favorable interaction site for the SMG1**i** urea group (Gly3943 at the position of SMG1 Asn2356) (Fig. 2 Supp. 4 F). In addition, DNA-PK PRD residue Lys4019 would sterically clash with the SMG1**i** urea group in the inactive conformation of the DNA-PK active site. In the case of ATR, the SMG1**i** sulfonamide site has diverged (Gly2385 at the position of SMG1 Pro2213) and residues in the N-lobe would result in a steric clash with the urea group of the inhibitor (Lys2329 and Asp2330 corresponding to SMG1 Leu2157 and Glu2158) (Fig. 2 Supp. 4 G). Finally, mTOR, DNA-PK and ATR active sites have a different relative orientation of the kinase N- and C-lobes as compared to SMG1, changing the overall chemical environment at the SMG1**i**-binding site (Fig. 2 C and Fig. 2 Supp. 4 F and G). We conclude that subtle differences in the SMG1 structure underpin the specificity of the SMG1**i** inhibitor.

### The SMG1 insertion domain contains a PRD that blocks substrate binding in the presence of SMG8

In the reconstruction of the SMG1**i**-bound SMG1-8-9 complex, we observed additional density near the end of the kinase domain, protruding from the position where the insertion domain is expected to start (Fig. 3 and Fig. 3 Supp. 1 A and B). While the lower resolution for this part of the map was not sufficient to unambiguously build an atomic model, this density clearly docks at the substrate-binding site of the kinase domain (Fig. 3 A, Fig. 3 Supp. 1 A, B and C). Superposition with the previous cryo-EM structure of SMG1-8-9 complex bound to a UPF1 peptide (Langer et al. 2020) revealed that the additional density in the current reconstruction is mutually exclusive with the position of a bound substrate (Fig. 3 I, Fig. 3 Supp. 1 A). This observation is in good agreement with the reported positions of the PRDs in structures of the autoinhibited forms of yeast Tel1^ATM^ and human DNA-PK (Jansma et al. 2020; Yates et al. 2020; Chen et al. 2021). We concluded that the SMG1 insertion domain contains a PRD that can directly occupy and block the substrate binding path in the kinase active site, analogous to other PIKKs. The SMG1 PRD would have to re-arrange upon substrate binding, as observed in the case of the inactive-to-active transition of DNA-PK (Chen et al. 2021).

**Figure 3:**
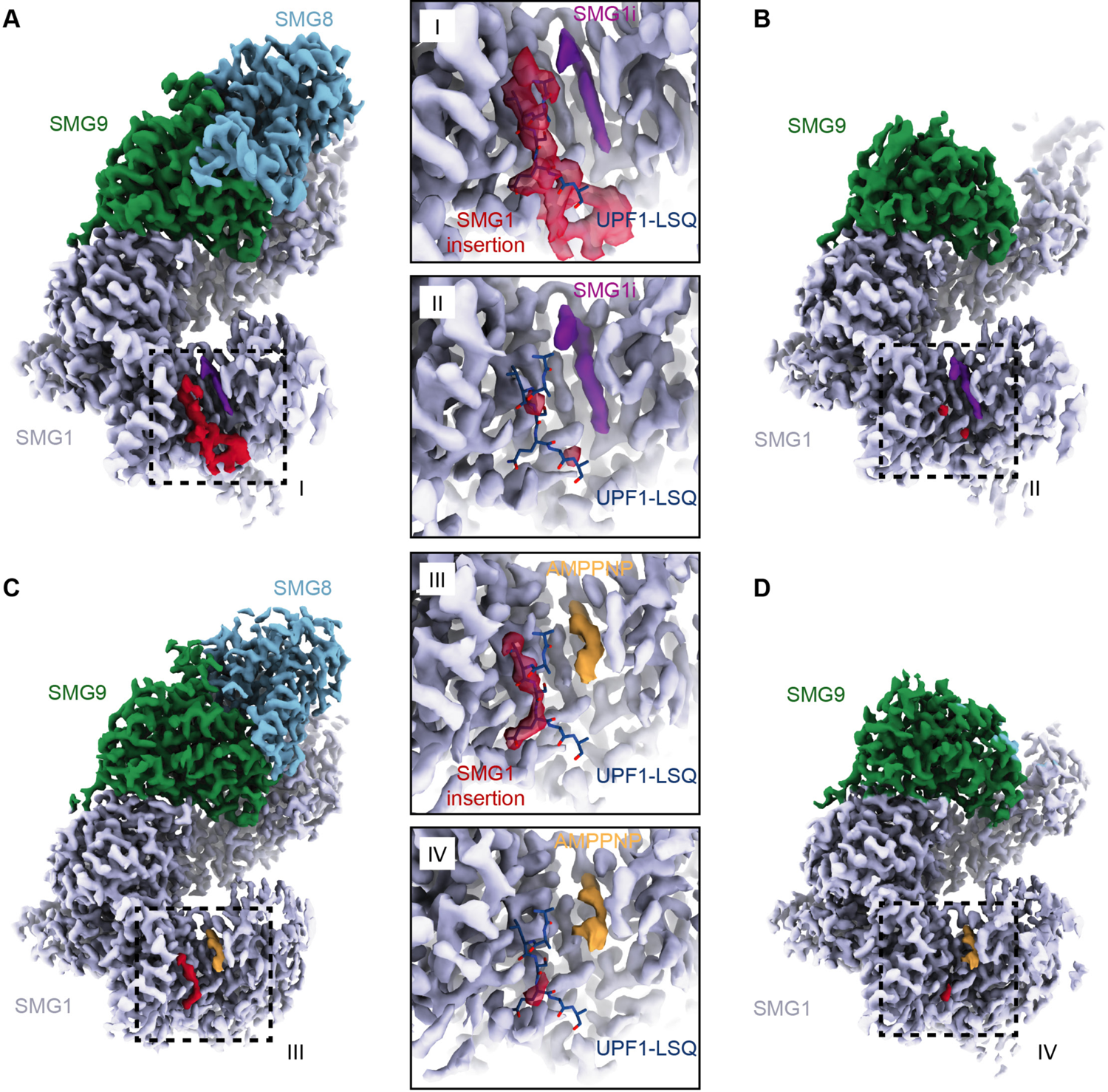
Structures of SMG1-8-9 and SMG1-9 complexes reveal that the SMG1 insertion domain can block the substrate-binding path in the presence of SMG8. A) Cryo-EM density of SMG1-8-9 bound to SMG1**i**. Density for the inhibitor is in magenta, the N-terminus of the SMG1 insertion is in red, and all other parts as indicated. B) Cryo-EM density of SMG1-9 complex bound to the SMG1 inhibitor. Everything else as in A). C) Cryo-EM density of SMG1-8-9 bound to AMPPNP. Density for AMPPNP is in orange and extra density attributed to the SMG1 insertion is in red. D) as in C) but for the SMG1-9 complex bound to AMPPNP. Insets I to IV show close-ups of the indicated kinase active site densities superimposed with the model for the UPF1-LSQ substrate shown as blue sticks (PDB: 6z3r).

Within the cryo-EM data set collected on a SMG1-8-9 complex incubated with SMG1**i**, we isolated a subset of particles missing any density for SMG8, hence representing a SMG1**i**-bound SMG1-9 complex (Fig. 3 B, Fig. 2 Supp. 1 C, Fig. 3 Supp. 2). The corresponding 3.6 Å reconstruction not only showed a similar mode of interaction of SMG1**i**, but also several features distinct from SMG1 in isolation and the SMG1-8-9 ternary complex. Globally, the *α*-solenoid of SMG1 rearranges, consistent with the notion of a stepwise compaction of SMG1 upon binding first SMG9 and then SMG8 (Fig. 3 Supp 2 A and B) (Melero et al. 2014; Zhu et al. 2019). Locally, the mode of interaction between SMG9 and the SMG1 *α*-solenoid differs between the two complexes. In the binary SMG1-9 complex, the portion of SMG9 that interacts with the N-terminal HEAT repeat helices of SMG1 changes conformation, exploiting part of a binding site that was occupied by a short N-terminal segment of SMG8 in the ternary complex (Fig. 3 Supp. 2 C, D and E). Indeed, two hydrophobic residues of SMG9 (Leu456 and Leu457) take the place normally reserved for two hydrophobic residues of SMG8 (Leu351 and Leu352) (Fig. 3 Supp. 2 F and G). Hence, the conformation of the SMG9 segment observed in the SMG1-9 complex is incompatible with binding of SMG8 to this part of SMG1. Importantly, the reconstruction of SMG1-9 showed no ordered density at the active site for the PRD (Fig. 3 B and II, Fig. 3 Supp. 1 A), which is in contrast to the ordered PRD density visualized in the SMG1**i**-bound SMG1-8-9 complex.

Next, we asked whether the autoinhibitory state of the SMG1 insertion domain in the SMG1-8-9 complex is due specifically to the presence of the SMG inhibitor or whether it is a more general feature. To address this question, we reconstituted purified wild-type SMG1-8-9 with an excess of AMPPNP (instead of SMG1**i**) and subjected the sample to a similar single-particle cryo-EM analysis routine. We obtained maps for SMG1-8-9 and for SMG1-9 at overall resolutions of 2.8 Å and 3.1 Å, respectively (Fig. 3 C and D, Fig. 2 Supp. 1 and 3). Both reconstructions showed clear density for AMPPNP in the SMG1 active site. In the reconstruction of the ternary SMG1-8-9 complex, we observed an additional density at the substrate binding site, in the same position as that observed when in the presence of SMG1**i** (Fig. 3 III and IV). However, in the context of AMPPNP this additional density is less prominent and does not connect to the SMG1 kinase domain. It thus appears that the SMG1 inhibitor has a stabilizing effect on the autoinhibitory conformation of the SMG1 PRD. In the reconstruction of the binary SMG1-9 complex in the presence of AMPPNP, we observed no ordered density at the substrate-binding site, consistent with the reconstruction obtained in the presence of SMG1**i**. We concluded that the autoinhibitory conformation of the PRD in the insertion domain of SMG1 is connected to the presence of a nucleotide in the ATP-binding site and is stabilized by the presence of SMG8.

### The SMG1 insertion domain can block overall access to the kinase active site and interacts with the SMG8 C-terminus

The findings above suggested a possible role for SMG8 in stabilizing an autoinhibited state of SMG1. Indeed, previous work implicated SMG8 and, in particular, its C-terminal domain in the down-regulation of SMG1 activity (Zhu et al. 2019; Arias-Palomo et al. 2011; Deniaud et al. 2015; Yamashita et al. 2009). SMG8 has a modular domain organization: the N-terminal G-domain is followed by a helical stalk that protrudes into solvent with well-defined density, but the remaining C-terminal region (amounting to about 45% of the molecule) is flexible and poorly resolved in the published cryo-EM studies (Gat et al. 2019; Zhu et al. 2019; Langer et al. 2020). By further processing of the SMG1**i**-bound SMG1-8-9 data, we could achieve improved density for the C-terminal domain of SMG8, showing a knob-like feature directly connected to the end of the stalk and in turn connecting to a larger globular mass (Fig. 4 A). Concurrently, we observed an additional density extending from the location of the PRD that we attributed to the SMG1 insertion domain (see below). On one side, this density wraps along the catalytic module of SMG1, occludes the access to the kinase active site (Fig. 4 A) and reaches towards the IP_6_-binding site (Fig 4 A and C). On the other side, the density projects into solvent and approaches the globular SMG8 C-terminal region. Although the low resolution of this part of the map did not allow model building, the close proximity suggested a physical contact between the SMG1 insertion domain and the SMG8 C-terminal domain. We proceeded to test the interplay between these two parts of the SMG1-8-9 complex.

**Figure 4:**
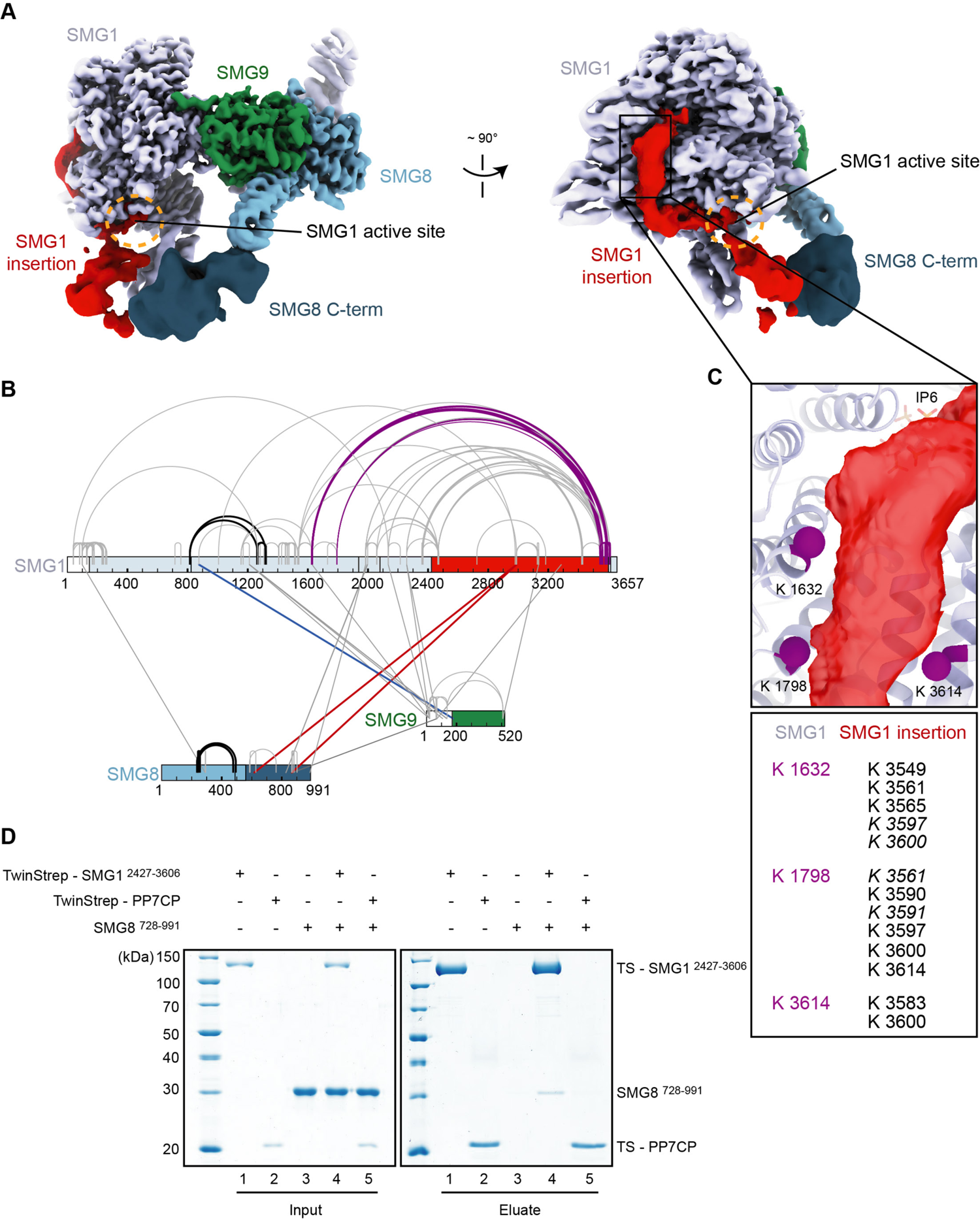
The SMG1 insertion domain can block overall access to the kinase active site. A) Cryo-EM map after 3D variability analysis filtered by resolution and segmented. Two different views displaying extra density for SMG8 C-terminus (dark blue) and SMG1 insertion domain. The location of the SMG1 kinase active site is indicated by an orange circle. B) Inter and intra cross-links of the SMG1-8-9 kinase complex. Proteins are colored as in A). Intra-links shown in Fig. 4 Supp. 1 B) and C) are in black, the inter-link shown in Fig. 4 Supp. 1 D) is in blue and inter-links between SMG1 insertion and SMG1 C-terminus are in red. C) Zoom-in highlighting positions of Lys residues (magenta spheres) crosslinking to the C-terminus of the SMG1 insertion domain. A table listing the cross-linked residues is shown below. Cross-links found in only one of two samples are in italics. D) Coomassie-stained SDS-PAGE analysis of pull-down experiment showing an interaction between SMG1 insertion domain (SMG1 ^2427-3606^) and SMG8 C-terminus (SMG8 ^728-991^).

We first used cross-linking mass spectrometry (XL-MS) analysis (Fig. 4 B, Fig. 4 Supp. 1 and Supp. 2). Upon treatment of purified SMG1-8-9 with Bis(sulfosuccinimidyl)suberate (BS^3^), we detected intra and inter cross-links consistent with parts of the complex for which an atomic model was available and measuring well within the expected range of distance for BS^3^ cross-links (Fig. 4 B, Fig. 4 Supp. 1 and Fig. 4 Supp. 2). We also observed reproducible intra cross-links between the end of the SMG1 insertion and the catalytic module, in particular the FAT (residues 823 to 1938) and kinase (residues 2080 to 2422) domains (Fig. 4 B and C). These cross-links mapped in close proximity to the aforementioned density extending from the PRD. We thus attribute this density to the C-terminus of the SMG1 insertion (Fig. 4 A, B and C). Importantly, we detected inter cross-links between the SMG1 insertion domain and the SMG8 C-terminal domain (Fig. 4 B), consistent with the proximity of the EM densities (Fig. 4 A) and previously reported XL-MS data (Deniaud et al. 2015).

Next, we tested whether these parts of the complex can interact directly in biochemical assays with recombinant proteins. We purified SMG1 insertion domain (SMG1 residues 2427-3606) with a N-terminal TwinStrep-tag from transiently transfected HEK293T cells and SMG8 C-terminus (SMG8 residues 728-991) from *E.coli*. Since the structural analysis is indicative of a dynamic or weak contact, we performed pull-down assays in low-salt conditions. We observed that the SMG1 insertion domain indeed co-precipitated the SMG8 C-terminus in these conditions (Fig. 4 D). As a control, an unrelated TwinStrep-tagged protein (PP7 coat protein) did not pull down SMG8 C-terminus under the same conditions (Fig. 4 D). As anticipated, increasing the salt concentration had a detrimental effect on the SMG8 C-terminus – SMG1 insertion co-precipitation (Fig. 4 Supp. 3 A).

Taken together, these experiments indicate the existence not just of proximity but of a direct physical link between the SMG1 insertion domain and SMG8 C-terminus, rationalizing the observed dependency of SMG1 autoinhibition on the presence of SMG8 (Fig. 3).

## Conclusions

In this manuscript, we reported a reconstruction of the SMG1-8-9 complex bound to a SMG1 inhibitor. We showed that this compound binds to the ATP-binding site within the kinase active site. By comparison with structural data of related kinases, we identified two functional groups within the small molecule that possibly confer specificity to SMG1. Using the SMG1-related mTOR kinase as an example, we confirmed the specific action of the inhibitor biochemically.

In addition to the SMG1 inhibitor, we observed density for the N-terminal part of the SMG1 insertion domain in the SMG1-8-9 reconstruction. This part of the insertion domain acts as a PRD and occupied the substrate binding path within the SMG1 active site. Consistent with data for other PIKKs, a PRD within the SMG1 insertion domain can therefore restrict access to the kinase active site.

The structure of a SMG1**i**-bound SMG1-9 complex reconstructed from the same data set as SMG1**i**-bound SMG1-8-9 did not show ordered density for the SMG1 PRD in the SMG1 active site. This suggested that an interaction between the SMG1 insertion and SMG8 was important for stabilizing the autoinhibited state of the complex. Consistently, our biochemical analysis indicated a direct physical link between parts of the SMG1 insertion domain and the C-terminal domain of SMG8. While autoinhibition was not observed in previously published apo-structures, our reconstructions of SMG1-9 and SMG1-8-9 bound to AMPPNP showed that blockage of the substrate binding path was also possible when the SMG1-8-9 complex active site was bound to a nucleotide. Accordingly, the described autoinhibition was not a phenomenon limited to the inhibitor-bound structures. We conclude that at least two layers of regulation of SMG1 kinase activity exist: First, access to the SMG1 active site can be restricted in *cis* by a PRD within the SMG1 insertion domain. Second, this autoinhibition is stabilized in *trans* by the SMG8 C-terminus.

How is the autoinhibited complex activated? The position of the SMG1 insertion domain in the inhibitor-bound SMG1-8-9 complex reconstruction raised the possibility that the autoinhibited complex might not be able to interact with its substrate, UPF1 (Fig. 3 I, III and Fig. 4 A). To test this hypothesis, we incubated TwinStrep-tagged SMG1-8-9 with increasing concentrations of SMG1**i** before adding UPF1 and performed a pull-down experiment (Fig. 4 Supp. 3 B). The efficiency of UPF1 co-precipitation was unaffected by the presence of SMG1**i**, suggesting that either UPF1 can overcome blockage of the SMG1 active site by the SMG1 insertion domain or has secondary binding sites independent of the observed SMG1 autoinhibition. Indeed, superposition of the autoinhibited complex reconstruction with a previously reported negative-stain map of a cross-linked UPF1-bound SMG1-8-9 complex raised the possibility that UPF1-binding and SMG1 autoinhibition might be able to occur in parallel (Fig. 4 Supp. 3 C).

Whether SMG8 or a SMG8-9 complex dissociates from SMG1 at any point in the kinases’ catalytic cycle during NMD *in vivo* is an outstanding question. Early biochemical experiments have suggested that SMG8 plays a crucial role in recruiting SMG1 complex to the NMD-competent messenger ribonucleoprotein particle (Yamashita et al. 2009). Given that the SMG1-8-9 complex actively phosphorylated UPF1 in *in vitro* experiments using purified proteins and our observation that SMG1**i** binding did not impair interaction of SMG1 with UPF1, it is tempting to speculate that UPF1 itself might be sufficient to overcome the effect of SMG8 on stabilizing SMG1 autoinhibition within a SMG1-8-9 complex. In this model, nucleotide binding and kinase activation could occur in two distinct steps during NMD: Upon ATP binding, the kinase complex would adopt an autoinhibited conformation and only be licensed for specific phosphorylation of its target by the substrate itself (Fig. 5). In addition, autophosphorylation of the kinase complex may play a crucial role in kinase activation.

**Figure 5:**
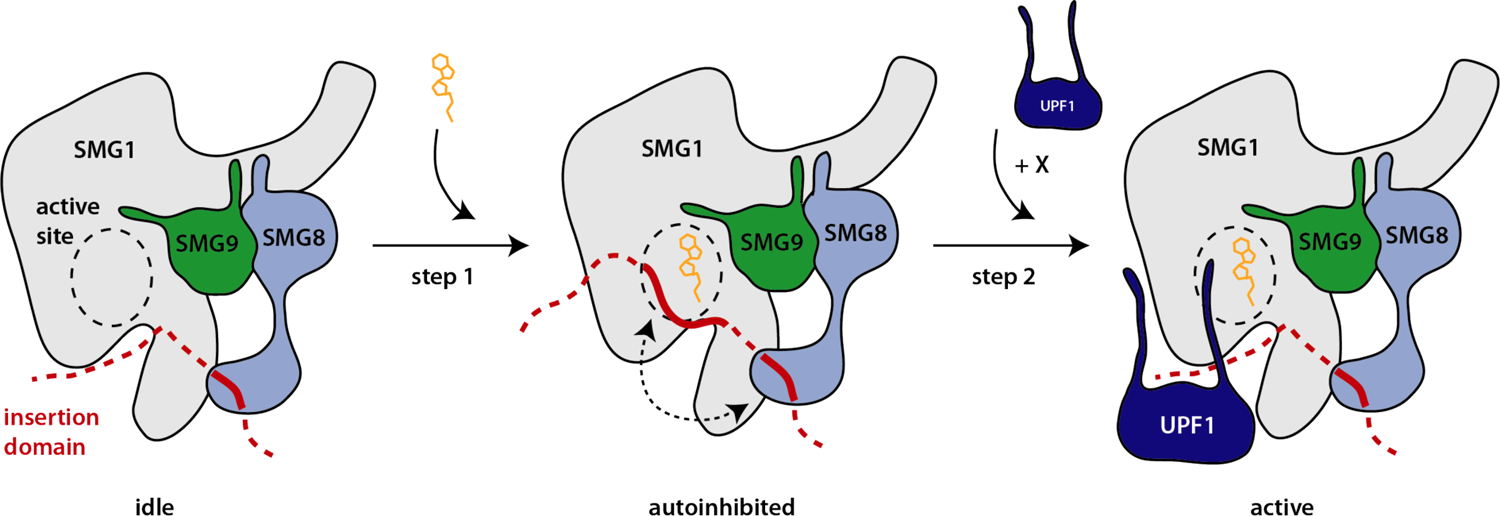
Hypothetical model of SMG1 kinase regulation. Structural and biochemical data suggest different layers of regulation on SMG1 kinase activity. Upon ATP binding (orange), the SMG1-8-9 complex adopts an autoinhibited conformation (step 1) - mediated by the concerted action of the SMG1 insertion domain in *cis* and the SMG8 C-terminus in *trans* (dotted arrow). The presence of the correct substrate and likely other factors/cues (indicated by X) then trigger the release of the autoinhibition and activity of the complex towards UPF1 (step 2).

Taken together, our structures will help to improve the design of selective PIKK active site inhibitors, e.g. specific for SMG1. Our data suggest a unifying structural model for concerted action of the SMG1 insertion domain in *cis* and SMG8 in *trans* to tune SMG1 kinase activity, shedding light on the intricate regulation of this kinase complex in metazoan nonsense-mediated mRNA decay.

## Materials and Methods

### Key Resources Table

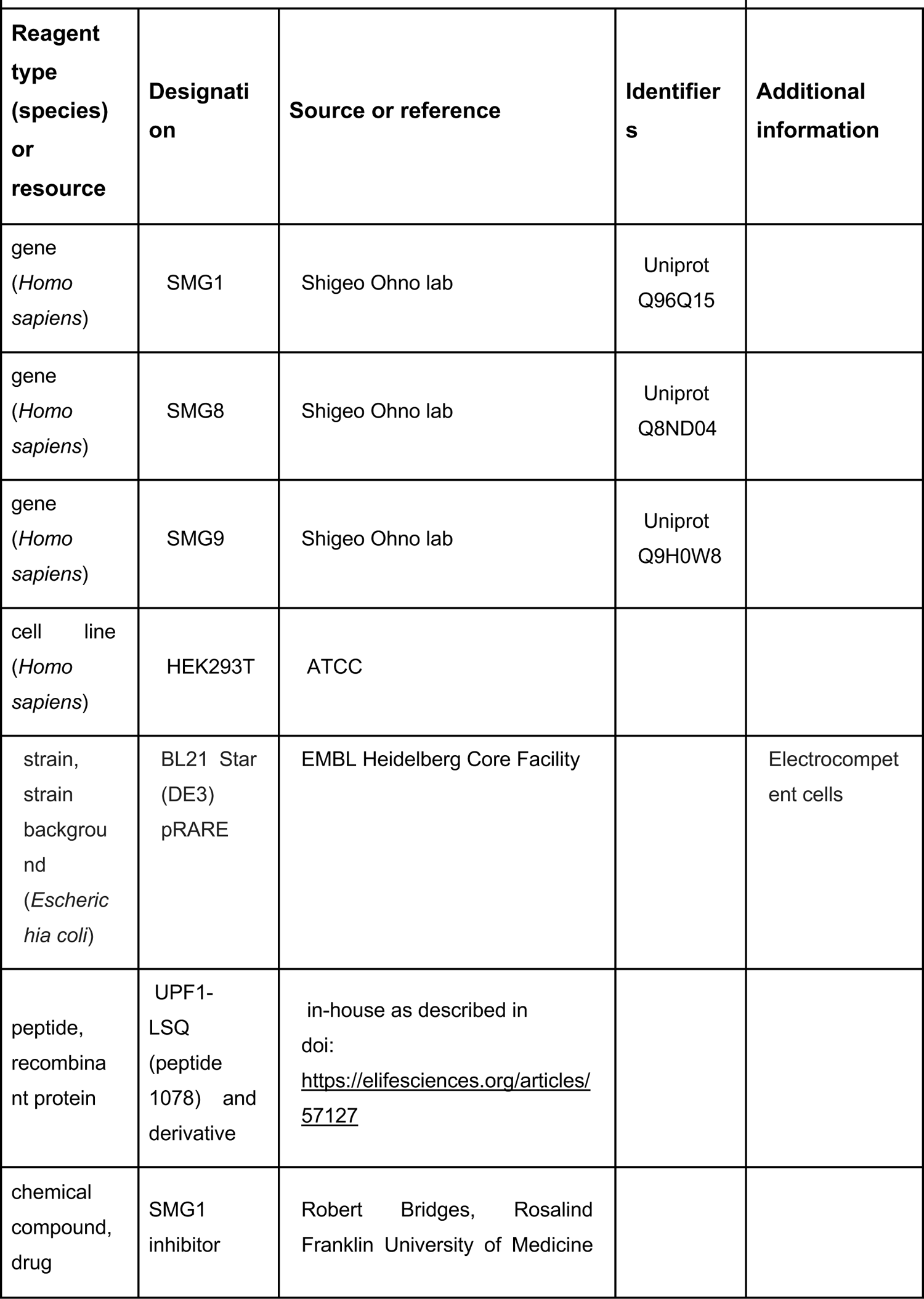
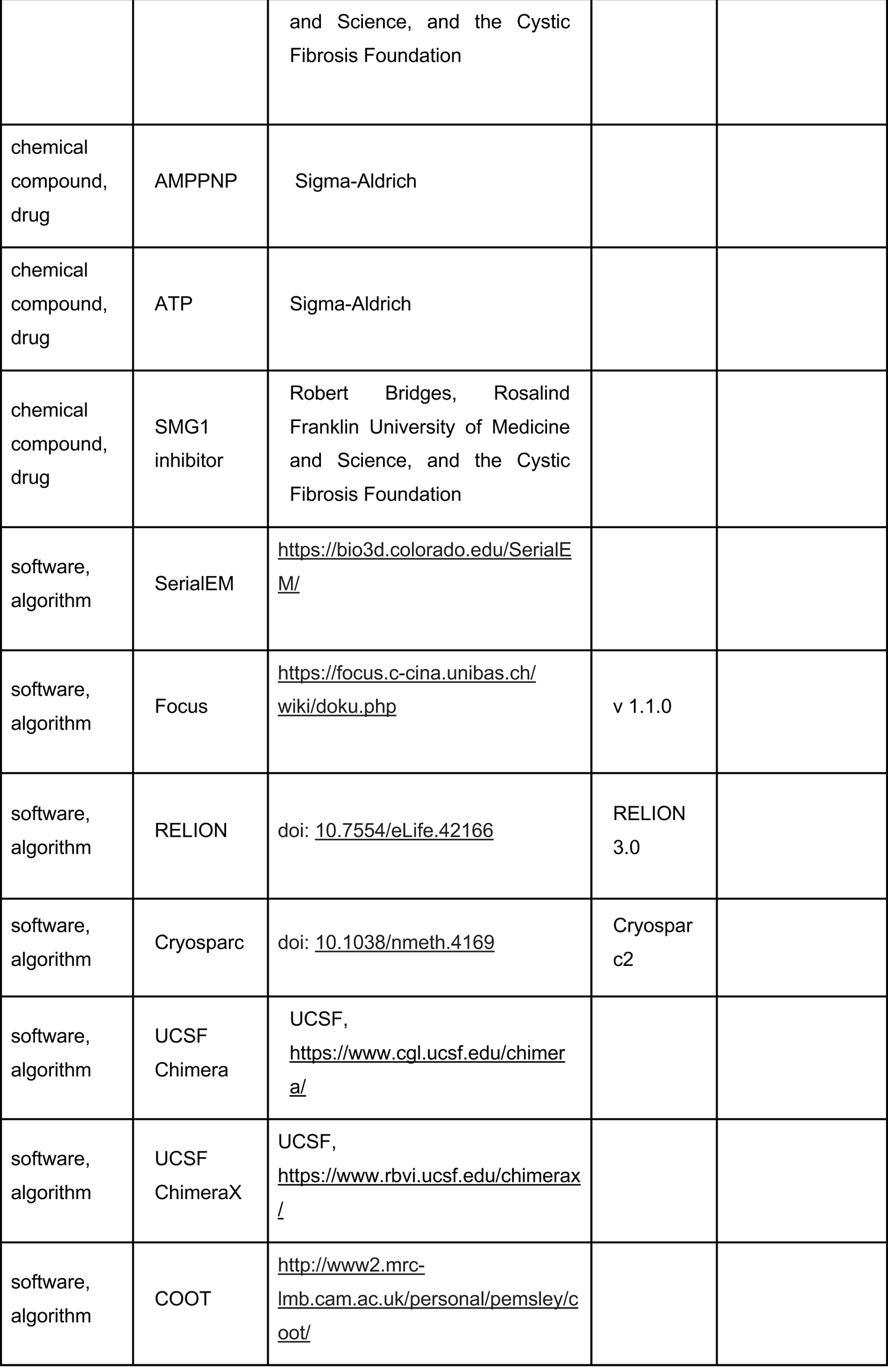
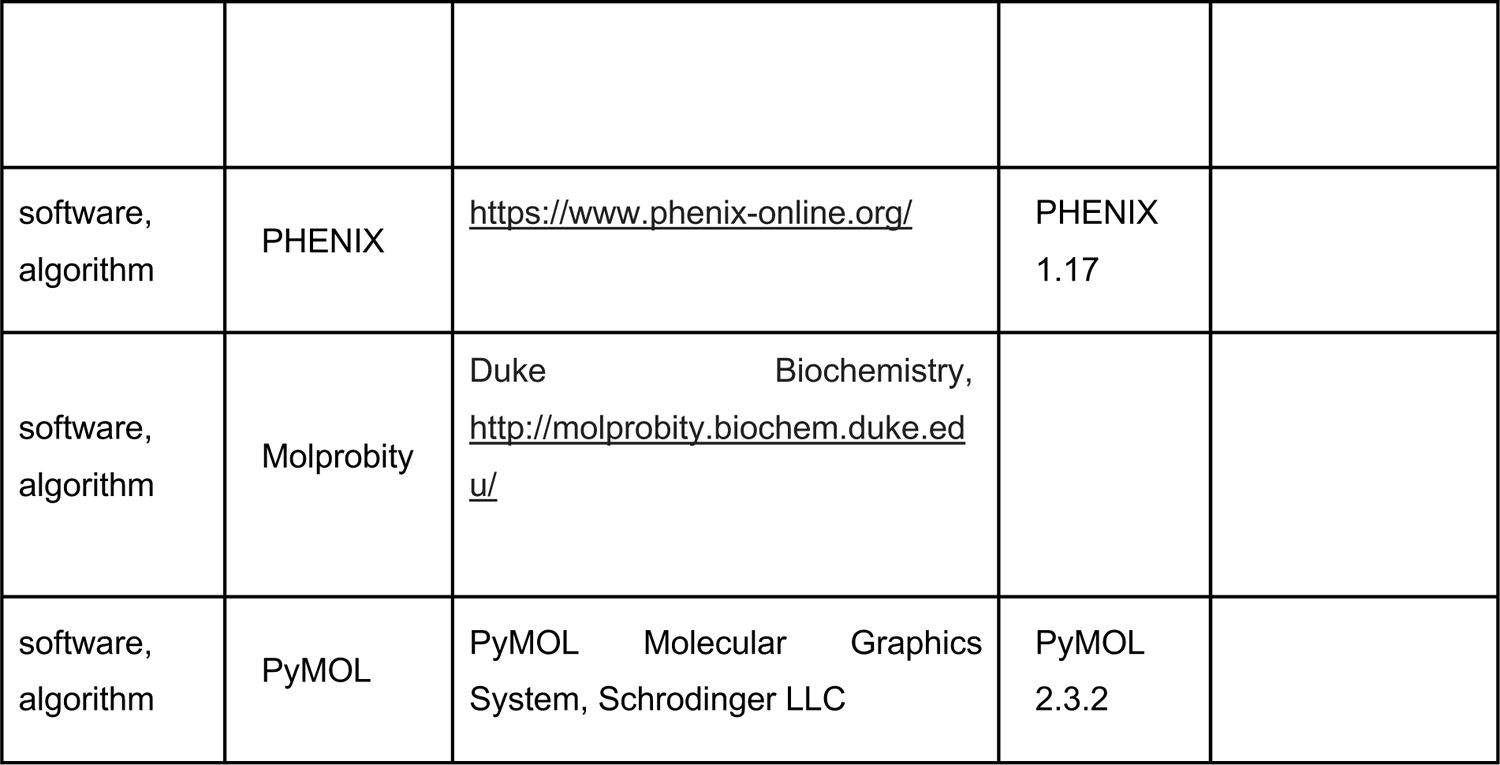

### Protein expression and purification

TwinStrep-tagged SMG1-8-9 complex was expressed and purified as described before (Gat et al. 2019; Langer et al. 2020). N-terminally TwinStrep-tagged SMG1 insertion domain (SMG1 ^2427-3606^) was expressed and purified essentially identical to the SMG1-8-9 full-length complex, with the exception that lysis and affinity purification was carried out in a buffer based on 2xPBS.

Untagged full-length UPF1 expressed in *Escherichia coli* was purified as reported previously (Langer et al. 2020). N-terminally TwinStrep-tagged full length UPF1 was expressed using a stable pool of HEK293T cells as described for SMG1-8-9. Approximately 400 x 10^6^ cells were lysed with a Dounce homogenizer in buffer containing 1xPBS, 5 mM MgCl_2_, 1 µM ZnCl_2_, 10 % (v/v) glycerol, 1 mM DTT supplemented for lysis with Benzonase and DNase I as well as EDTA-free cOmplete Protease Inhibitor Cocktail (Roche). The lysate was cleared by centrifugation at 25,000 rpm for 30 min at 10°C. The supernatant was filtered and applied to a StrepTrap HP column (Sigma-Aldrich). After passing 25 column volumes of lysis buffer without supplements, the column was further washed with 25 column volumes of buffer Hep A (20 mM HEPES/NaOH pH 7.4, 85 mM KCl, 2 mM MgCl_2_, 1 µM ZnCl_2,_ 10 % (v/v) glycerol, 1 mM DTT) and bound protein was eluted directly onto a HiTrap Heparin column (GE Healthcare) using buffer Hep A supplemented with 2.5 mM Desthiobiotin. After washing the Heparin column with 10 column volumes of buffer Hep A, bound protein was eluted using a gradient increasing the salt concentration to 500 mM KCl over 100 mL while collecting fractions (AEKTA prime FPLC system, GE Healthcare). Fractions were analyzed by SDS-PAGE and those containing pure full-length TwinStrep-UPF1 were collected, concentrated and flash frozen in liquid nitrogen until further usage.

The SMG8 C-terminus (SMG8 ^728-991^) was N-terminally fused to a 6xHis-SUMO tag and expressed in *E.coli* BL21 STAR (DE3) pRARE cells. Bacteria were grown in TB medium at 37°C shaking at 180 rpm. At an OD of 2, cultures were cooled down to 18°C and overnight protein expression was induced by adding IPTG. After harvesting (6,000 rpm, 10 min), bacteria were lysed by sonication in a buffer containing 20 mM Tris-HCl pH 7.5, 400 mM NaCl, 10 mM Imidazole pH 7.5, 1mM ß-mercaptoethanol, Benzonase, DNase I and 1 mM PMSF. After centrifugation (25,000 rpm, 30 min, 10°C) and filtration, the cleared lysate was applied to a HisTrap FF column (Cytiva) and the column was washed with 30 column volumes wash buffer (20 mM Tris-HCl pH 7.5, 400 mM NaCl, 40 mM Imidazole pH 7.5, 1mM ß-mercaptoethanol). Bound protein was eluted with wash buffer supplemented with 340 mM Imidazole pH 7.5 and the elution was combined with His-tagged SUMO protease for cleavage of the N-terminal tag and dialyzed overnight against 20 mM Tris-HCl pH 7.5, 400 mM NaCl, 1mM ß-mercaptoethanol. The dialyzed, protease treated sample was passed again over a 5 mL HisTrap FF column to remove cleaved tags and protease, the flow-through was diluted to 150 mM NaCl and 10 % (v/v) glycerol and applied to a HiTrap SP HP column (Cytiva). After washing with 10 column volumes of 20 mM Tris-Cl pH 7.5, 150 mM NaCl, 10 % (v/v) glycerol, and 1 mM ß-mercaptoethanol, proteins were eluted using a gradient increasing the salt concentration to 500 mM NaCl. The peak fractions containing SMG8 ^728-991^ were pooled, concentrated and injected onto a Superdex 75 10/300 GL (Cytiva) equilibrated with 20 mM HEPES/NaOH pH 7.4, 200 mM NaCl, and 1mM DTT. Again, the peak fraction containing SMG8 ^728-991^ was pooled and concentrated and flash frozen using liquid nitrogen. mTOR^ΔN^-LST8 and GST-AKT1 ^450-480^ were prepared as described before (Gat et al. 2019).

### Kinase assays

The mass spectrometry-based kinase assays were carried out as previously reported (Langer et al. 2020). For experiments involving SMG1**i**, the assembled reactions were incubated in the absence of ATP for 30 min at 4°C. All reactions involving SMG1**i** were performed with equal amounts of DMSO present.

Kinase assay based on the use of γ-^32^P-labelled ATP were carried out as reported. For all assays, 100 nM of kinase complex and 1 μM of substrate were used. All reactions were supplemented with equal amounts of DMSO and incubated without ATP for 30 min at 4°C before starting the experiment. After 30 min at 30°C, phosphorylation reactions were stopped by adding SDS-containing sample buffer and samples were analyzed using a SDS-polyacrylamide gel supplemented with 2,2,2-trichlorethanol (TCE, Acros Organics) for stain-free visualization (Ladner et al. 2004). The TCE-tryptophan reaction was started by exposing the protein gel to UV light for 2 min and proteins were visualized and quantified using a Bio-Rad gel imaging system and the Image Lab software (version 6.1, Bio-Rad) or by Coomassie-staining. Subsequently the gel was washed several times with water to remove residual γ-^32^P-labelled ATP and phosphoproteins were detected using autoradiography and a Amersham Typhoon RGB biomolecular imager (GE Healthcare). Phosphoprotein signal (P-UPF1 or P-mTOR^ΔN^) was quantified by densitometry using Fiji and normalized to the total amount of protein determined using TCE. Independent triplicates of each condition were normalized to the DMSO-only sample and plotted using Prism (GraphPad).

### Cross-linking mass spectrometry

For cross-linking mass spectrometry, 1 μM of SMG1-8-9 complex was incubated with 0.5 mM BS^3^ for 30 min on ice in a buffer containing 1xPBS, 5 mM MgCl_2_, and 1 mM DTT. The reaction was quenched by adding 40 mM Tris-HCl pH 7.9 and incubating for 20 min on ice. The sample was spun for 15 min at 18,000 g. For denaturation of the crosslinked proteins, 4 M Urea and 50 mM Tris was added to the supernatant and the samples were sonicated using a Bioruptor Plus sonication system (Diogenode) for 10x 30 sec at high intensity. For reduction and alkylation of the proteins, 40 mM 2-cloroacetamide (CAA, Sigma-Aldrich) and 10 mM tris(2-carboxyethyl)phosphine (TCEP; Thermo Fisher Scientific) were added. After incubation for 20 min at 37 °C, the samples were diluted 1:2 with MS grade water (VWR). Proteins were digested overnight at 37 °C by addition of 1 µg of trypsin (Promega). Thereafter, the solution was acidified with trifluoroacetic acid (TFA; Merck) to a final concentration of 1%, followed by desalting of the peptides using Sep-Pak C18 1cc vacuum cartridges (Waters). The elution was vacuum dried.

Enriched peptides were loaded onto a 30-cm analytical column (inner diameter: 75 microns; packed in-house with ReproSil-Pur C18-AQ 1.9-micron beads, Dr. Maisch GmbH) by the Thermo Easy-nLC 1000 (Thermo Fisher Scientific) with buffer A (0.1% (v/v) Formic acid) at 400 nl/min. The analytical column was heated to 60 °C. Using the nanoelectrospray interface, eluting peptides were sprayed into the benchtop Orbitrap Q Exactive HF (Thermo Fisher Scientific)(Hosp et al. 2015). As gradient, the following steps were programmed with increasing addition of buffer B (80% Acetonitrile, 0.1% Formic acid): linear increase from 8 to 30% over 60 minutes, followed by a linear increase to 60% over 5 minutes, a linear increase to 95% over the next 5 minutes, and finally maintenance at 95% for another 5 minutes. The mass spectrometer was operated in data-dependent mode with survey scans from m/z 300 to 1650 Th (resolution of 60k at m/z = 200 Th), and up to 15 of the most abundant precursors were selected and fragmented using stepped Higher-energy C-trap Dissociation (HCD with a normalized collision energy of value of 19, 27, 35). The MS2 spectra were recorded with dynamic m/z range (resolution of 30k at m/z = 200 Th). AGC target for MS1 and MS2 scans were set to 3 x 10^6^ and 10^5^, respectively, within a maximum injection time of 100 and 60 ms for the MS1 and MS2 scans, respectively. Charge state 2 was excluded from fragmentation to enrich the fragmentation scans for cross-linked peptide precursors.

The acquired raw data were processed using Proteome Discoverer (version 2.5.0.400) with the XlinkX/PD nodes integrated (Klykov et al. 2018). To identify the crosslinked peptide pairs, a database search was performed against a FASTA containing the sequences of the proteins under investigation. DSS was set as a crosslinker. Cysteine carbamidomethylation was set as fixed modification and methionine oxidation and protein N-term acetylation were set as dynamic modifications. Trypsin/P was specified as protease and up to two missed cleavages were allowed. Furthermore, identifications were only accepted with a minimal score of 40 and a minimal delta score of 4. Otherwise, standard settings were applied. Filtering at 1% false discovery rate (FDR) at peptide level was applied through the XlinkX Validator node with setting simple.

### Pull-down assays

Pull downs including the SMG8 C-terminus were performed using Strep-TactinXT Superflow beads (IBA) equilibrated with 20 mM HEPES/NaOH pH 7.4, 50 mM NaCl, 10% (w/v) glycerol, 1 mM DTT and 0.1% (w/v) NP-40 substitute (Fluka) unless stated otherwise. 1.5 μM TwinStrep-tagged protein was combined with 15 μM untagged protein in the buffer described above and incubated at 4°C for 30 min before adding equilibrated resin. The complete reaction was incubated for 1.5 hrs, beads were washed four times with 20x resin volume of buffer and bound protein eluted by adding buffer supplemented with 50mM Biotin.

To analyze the effect of SMG1**i** binding to SMG1 on the interaction between the SMG1-8-9 kinase complex and UPF1, 0.12 μM TwinStrep-tagged SMG1-8-9 was combined with DMSO or the respective amounts of SMG1**i** in buffer containing 1xPBS, 5 mM MgCl_2_, 1 mM DTT and 0.1% (v/v) NP-40. After incubation for 30 min at 4°C, 0.4 μM of untagged UPF1 was added and the reactions were combined with MagStrep “type3” XT beads (IBA). Samples were incubated for 30 min before washing four times with 20x resin volume of buffer and precipitation of bound proteins using SDS-containing sample buffer. For all pull-down assays, input and elution samples were analyzed by SDS-PAGE and stained with Coomassie.

### Cryo-EM sample preparation, data collection and data processing

Grids were prepared as described before (Langer et al. 2020), with the difference that SMG1-8-9 was incubated with either 4 μM SMG1i or 1 mM AMPPNP for 30 min on ice in 1xPBS, 5 mM MgCl_2_, and 1 mM DTT before adding 0.04 % (v/v) n-octyl-ß -D-glucoside and plunging using an ethane/propane mixture and a ThermoFisher FEI Vitrobot IV.

Cryo-EM data were essentially acquired as reported previously using a ThermoFisher FEI Titan Krios G3 microscope equipped with a post-column GIF (energy width 20 eV). The Gatan K3 camera was used in counting mode and data were acquired using SerialEM (Mastronarde 2005) and a beam-tilt based acquisition scheme. The nominal magnification during data collection for both data sets was 105.000x, corresponding to a pixel size of 0.8512 Å at the specimen level. The SMG1**i** sample was imaged with a total exposure of 89.32 e^-^/ Å^2^ evenly spread over 4 seconds and 40 frames. The AMPPNP sample was imaged with a total exposure of 60.99 e^-^/ Å^2^ evenly spread over 5.7 seconds and 40 frames at CDS mode. For both data collections, the target defocus ranged between −0.5 and −2.9 μm.

Movies were pre-processed on-the-fly using Focus (Biyani et al. 2017), while automatically discarding images of poor quality. Picked candidate particles were extracted in RELION 3.1 (Zivanov et al. 2018). After 2 rounds of reference-free 2D classification, particles were imported to CryoSPARC v2 (Punjani et al. 2017) for further processing in 2D and 3D. For details refer to Figure 2 - supplements 2 and 3. FSC curves were calculated using the 3D FSC online application (Tan et al. 2017).

### Model building

Model building was carried out using Coot (version 1.0) (Emsley et al. 2010) and iterative rounds of real-space refinement in the PHENIX software suit (Adams et al. 2011; Liebschner et al. 2019) and was based on our previously published models of SMG1-8-9 (PDB: 6syt, 6z3r). Geometric restraints for the SMG1**i** molecule were calculated with eLBOW (Moriarty, Grosse-Kunstleve, and Adams 2009). For details see Supplemental Table 1. Structure visualization and analysis was carried out using UCSF ChimeraX (version 1.2.5) (Goddard et al. 2018) and PyMOL (version 2.3.2).

## Acknowledgements

We are grateful to Robert Bridges, Rosalind Franklin University of Medicine and Science, and the Cystic Fibrosis Foundation for providing the SMG1 inhibitor. Daniel Bollschweiler and Tillman Schäfer at the MPIB cryo-EM facility for help with EM data collection and Barbara Steigenberger and Elisabeth Weyher at MPIB biochemistry core facility for carrying out mass spectrometry. We thank Christian Benda and J. Rajan Prabu for maintenance and development of computational infrastructure for EM data processing, Marcela Cueto for TwinStrep-PP7CP, Daniela Wartini for help with tissue culture and Courtney Long and members of the lab for help with the preparation of the manuscript. This work was supported by funding from the Max Planck Gesellschaft, the European Commission (ERC Advanced Investigator Grant EXORICO), and the German Research Foundation (DFG SFB1035, GRK1721, SFB/TRR 237) to E.C. and a Boehringer Ingelheim Fonds PhD fellowship to L.L.

---

**Figure 1 - Supp. 1:**
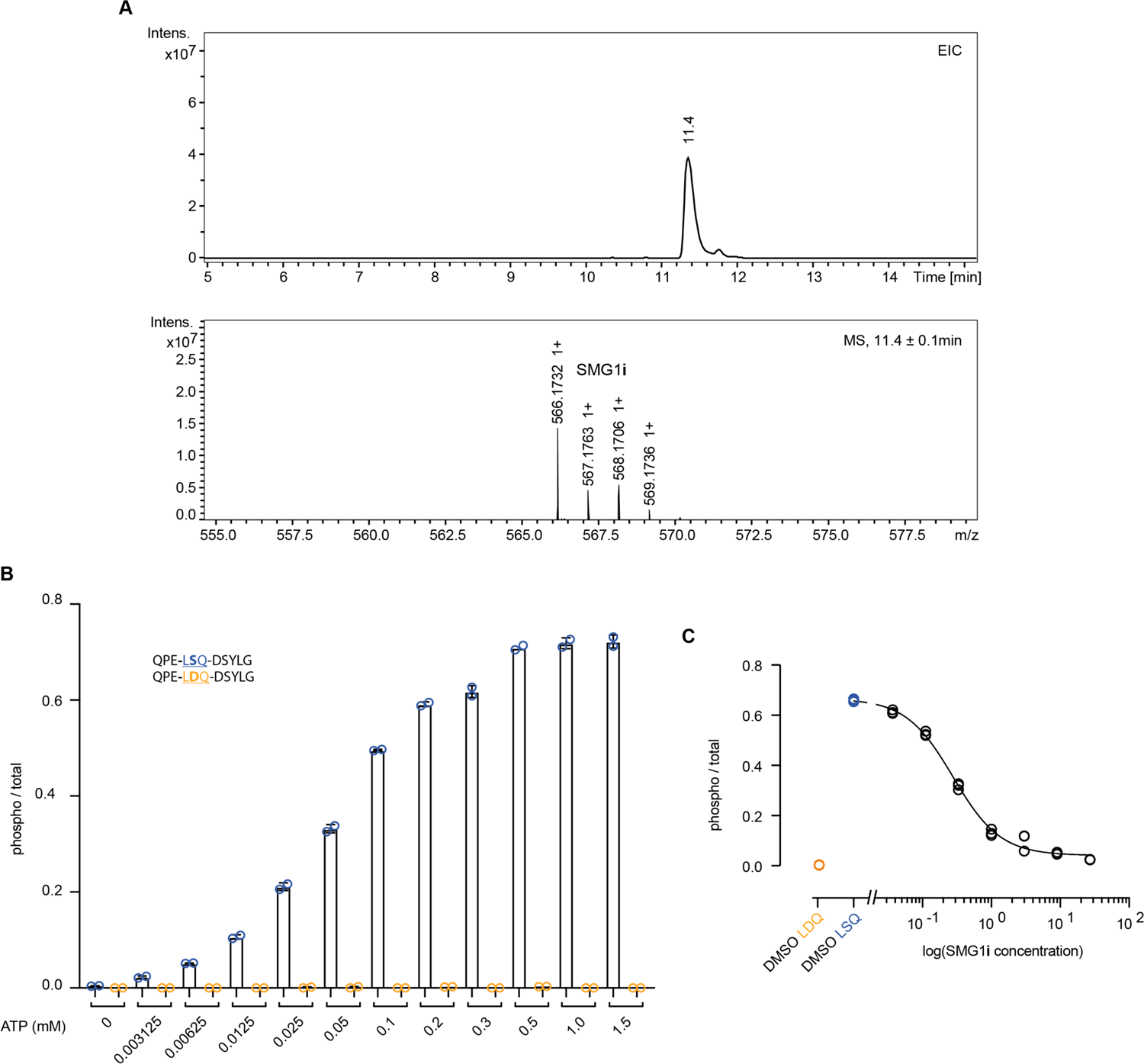
Characterization of SMG1 inhibitor. A) Liquid chromatography-mass spectrometry (LC-MS) experiment with the SMG1 inhibitor sample used throughout this study. The expected mass for SMG1**i** is 566.13 Da. Differences of +1 are caused by the varying composition of naturally occurring Carbon isotopes. B) Titration of ATP requirement in mass spectrometry-based phosphorylation assay using 500 nM SMG1-8-9 and the indicated UPF1-derived peptides as substrates. 0.75 mM ATP was chosen for later experiments. C) Dose-response curve for SMG1**i** in mass spectrometry-based phosphorylation assay (using data points shown in Figure 1B). Phosphorylation ratios of substrate (blue) and control peptide (orange) in the absence of SMG1**i** are shown. The control peptide was included for reference but was not used for fitting.

**Figure 1 - Supp. 2:**
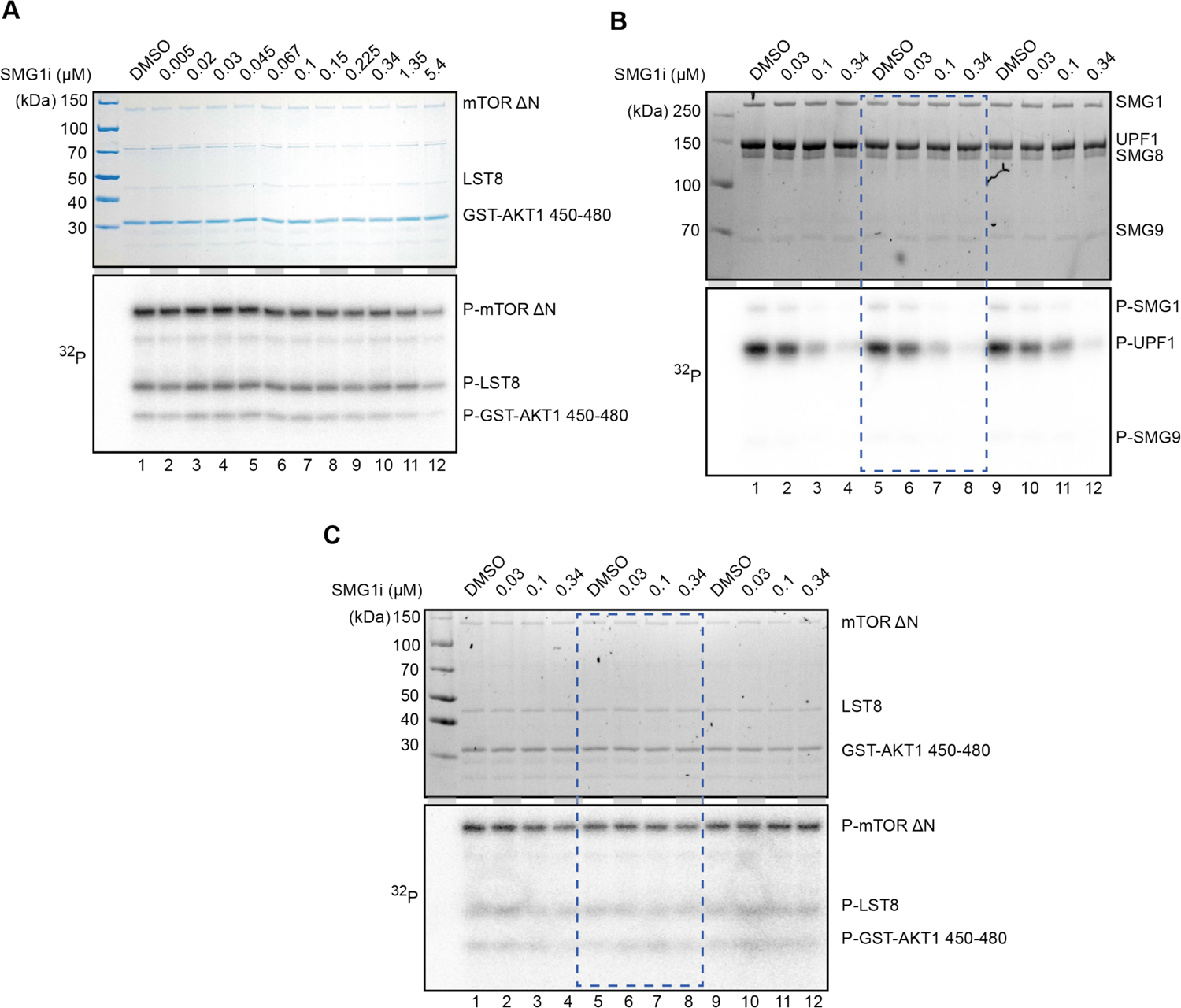
Titrations of SMG1 inhibitor with SMG1 and mTOR. A) Titration of SMG1**i** using a radioactivity-based phosphorylation assay with 30 nM mTOR-LST8 and GST-AKT1 as a substrate. The Coomassie-stained gel is shown on top and the radioactive signal on the bottom. B) and C) SMG1-8-9 or mTOR-LST8 radioactivity-based kinase assay with four selected concentrations of SMG1**i**. Assays were performed in triplicates and used for quantification. The blue rectangle indicates the replicate of the assays shown in Fig. 1.

**Figure 2 - Supp. 1:**
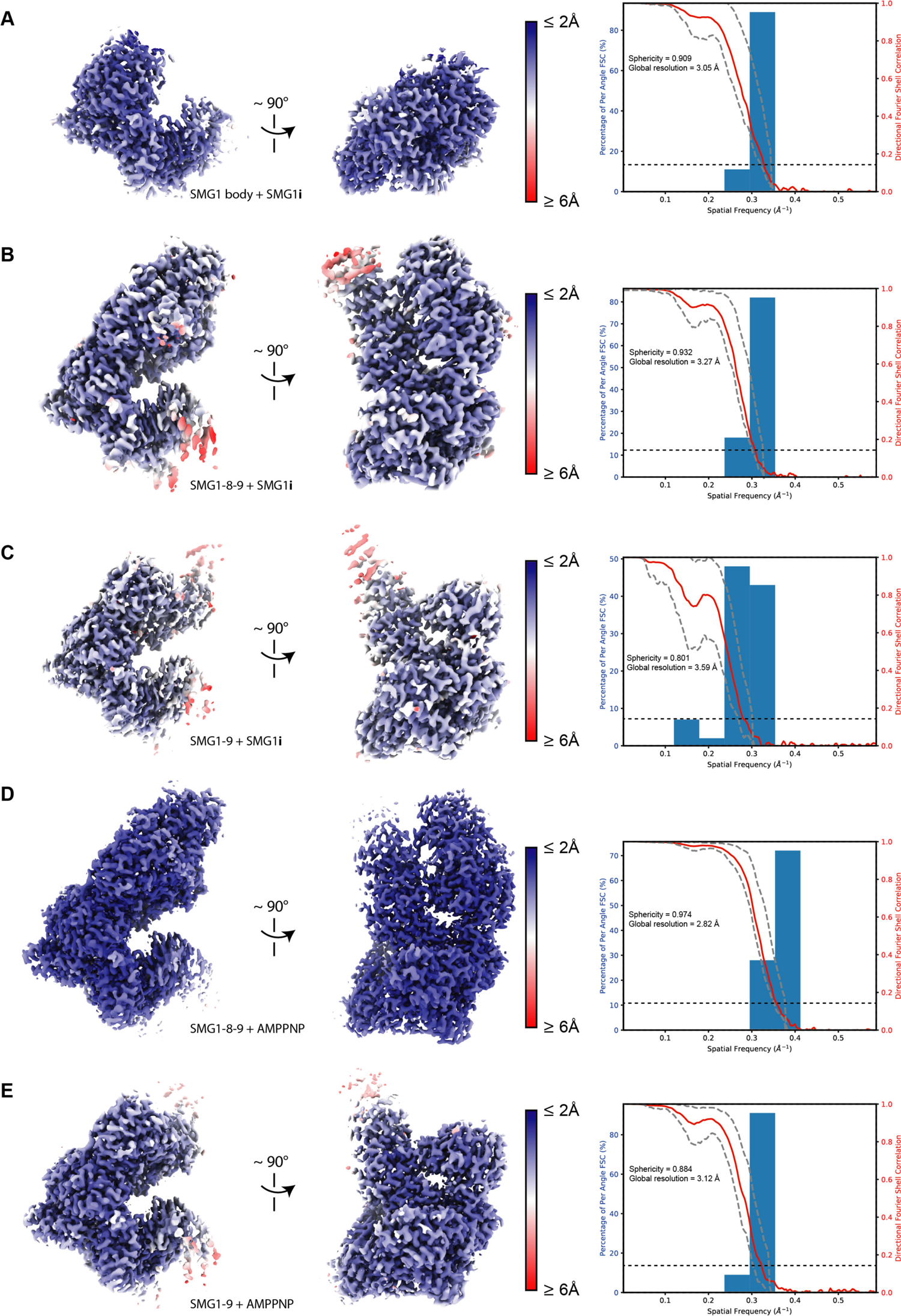
Resolution distribution and isotropy of SMG1-centered cryo-EM maps. Reconstructions used in this study for model building colored according to estimated local resolution shown in two different orientations. A three-dimensional FSC plot is included for each reconstruction (Tan et al. 2017): The red line represents the estimated global masked half map FSC. The resolutions according to the gold standard FSC cut off of 0.143 are indicated and shown as a black dashed line (Rosenthal and Henderson 2003). The spread of directional resolution values is defined as ± 1*σ* (dashed grey lines). Overall isotropy of the maps is indicated by the given sphericity values (out of 1). A) SMG1 body bound to SMG1**i** after focused refinement (EMDB XXXX). B) SMG1-8-9 bound to SMG1**i** (EMDB XXXX). C) SMG1-9 bound to SMG1**i** (EMDB XXXX). D) SMG1-8-9 bound to AMPPNP (EMDB XXXX). E) SMG1-9 bound to AMPPNP (EMDB XXXX).

**Figure 2 - Supp. 2:**
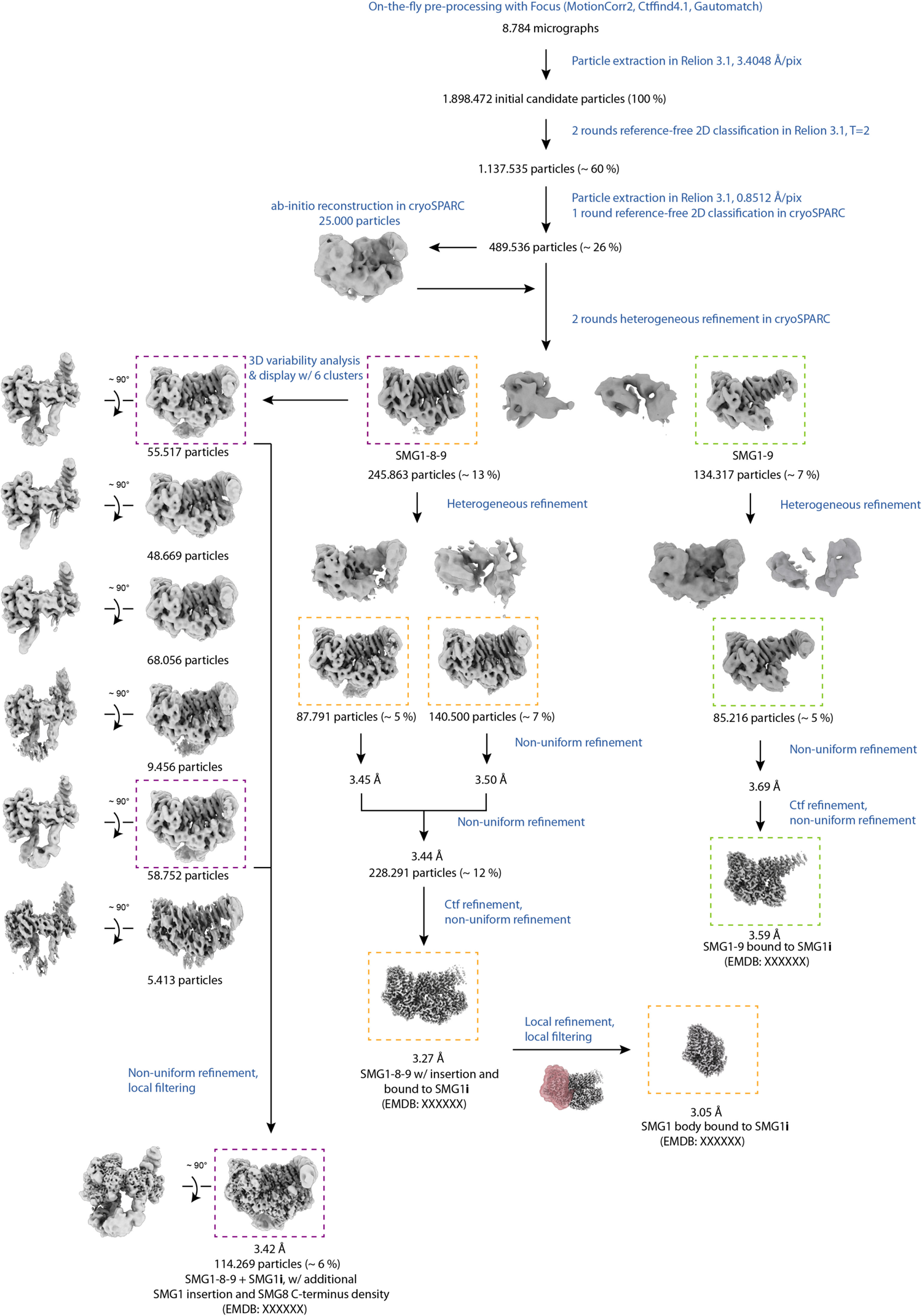
Cryo-EM data processing of SMG1i data set. Processing steps are indicated in blue; particle numbers and percentages with respect to initial candidate particles are shown for relevant classes. Colored, dashed rectangles indicate the different final reconstructions and the respective classes obtained from the data set collected in the presence of SMG1**i**.

**Figure 2 - Supp. 3:**
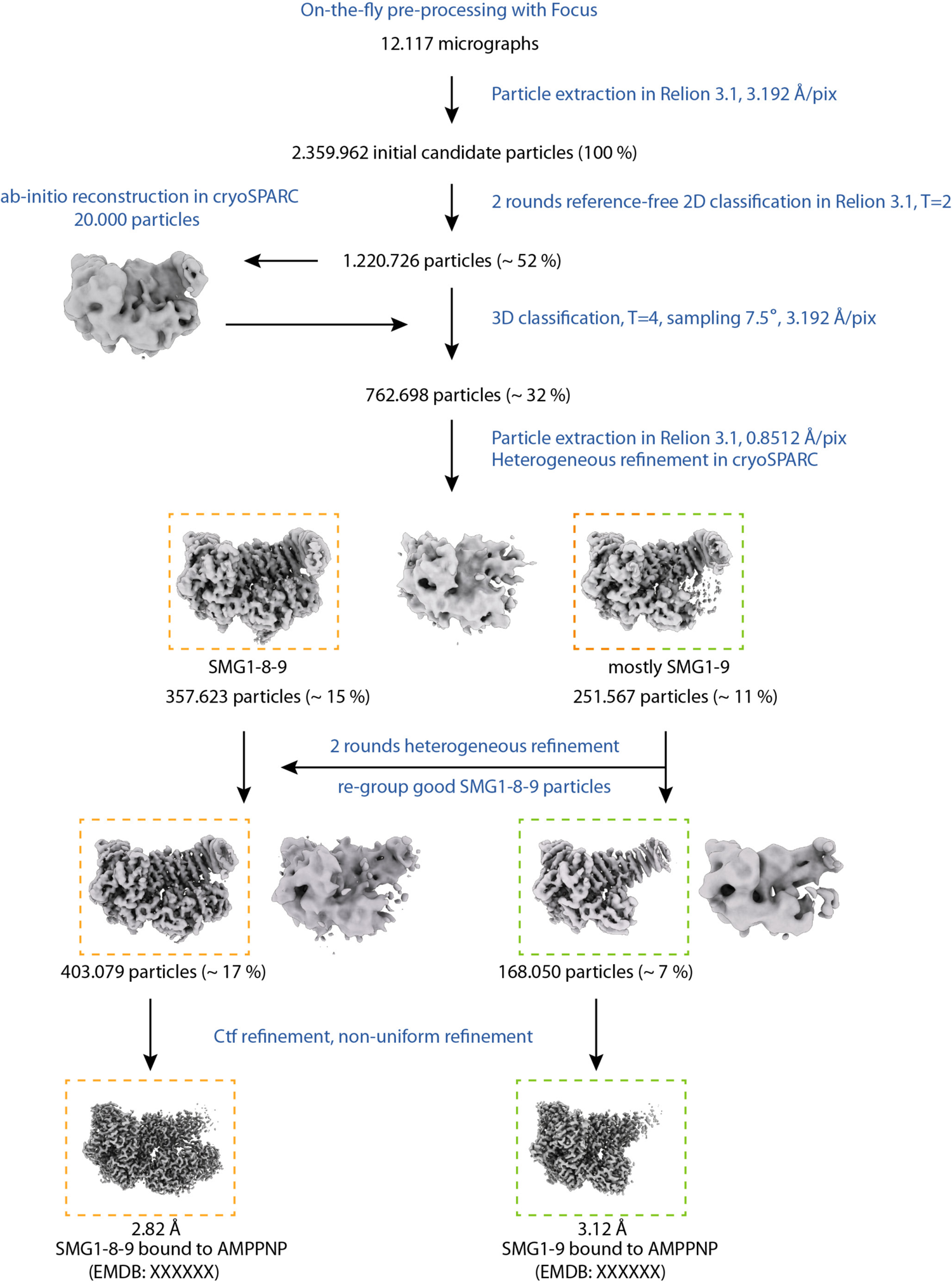
Cryo-EM data processing of AMPPNP data set. Processing steps are indicated in blue; particle numbers and percentages with respect to initial candidate particles are shown for relevant classes. Colored, dashed rectangles indicate the different final reconstructions and the respective classes obtained from the data set collected in the presence of AMPPNP.

**Figure 2 - Supp. 4:**
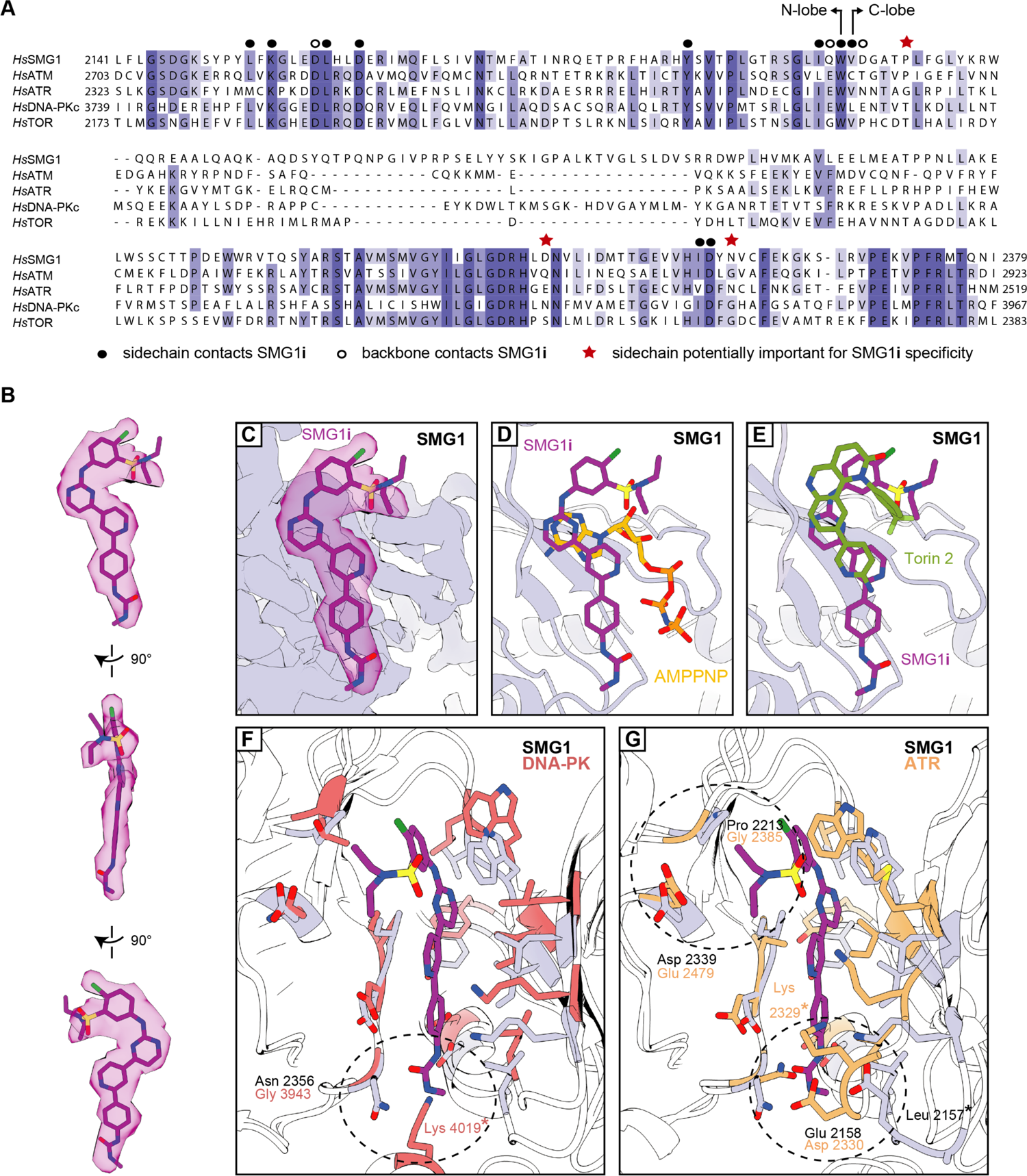
Further details of SMG1i binding and specificity. A) Multiple sequence alignment of parts of the kinase domains (N- and C-lobe indicated) belonging to the catalytically active members of the PIKK family with residues colored by identity. Residues of special interest are highlighted as indicated. B) Different views of the isolated cryo-EM density for SMG1**i** with the fitted model. C) Close-up side-view of cryo-EM density of SMG1-8-9 active site bound to SMG1**i**. The inhibitor model is shown and the corresponding segmented density is displayed in transparent magenta. D) Superposition of SMG1**i**-bound active site with AMPPNP-bound active site (PDB: 6z3r). Same view as in C). E) Superposition of SMG1**i**-bound active site with Torin 2-bound mTOR active site (PDB: 4jxp). Similar view as in C). F) Superposition of SMG1**i**-bound SMG1 with DNA-PK active site in the inactive conformation (PDB: 7k11). Figure prepared and labeled as in Fig. 2 C), with DNA-PK residues colored in red. The asterisk indicates a residue only visible in the DNA-PK structure due to structural rearrangements. G) Superposition of SMG1**i**-bound SMG1 with ATR active site (PDB: 5yz0). Figure prepared and labeled as in Fig. 2 C), with ATR residues colored in orange. An asterisk indicates corresponding residues in SMG1 and ATR that are separated due to conformational divergence.

**Figure 3 - Supp. 1:**
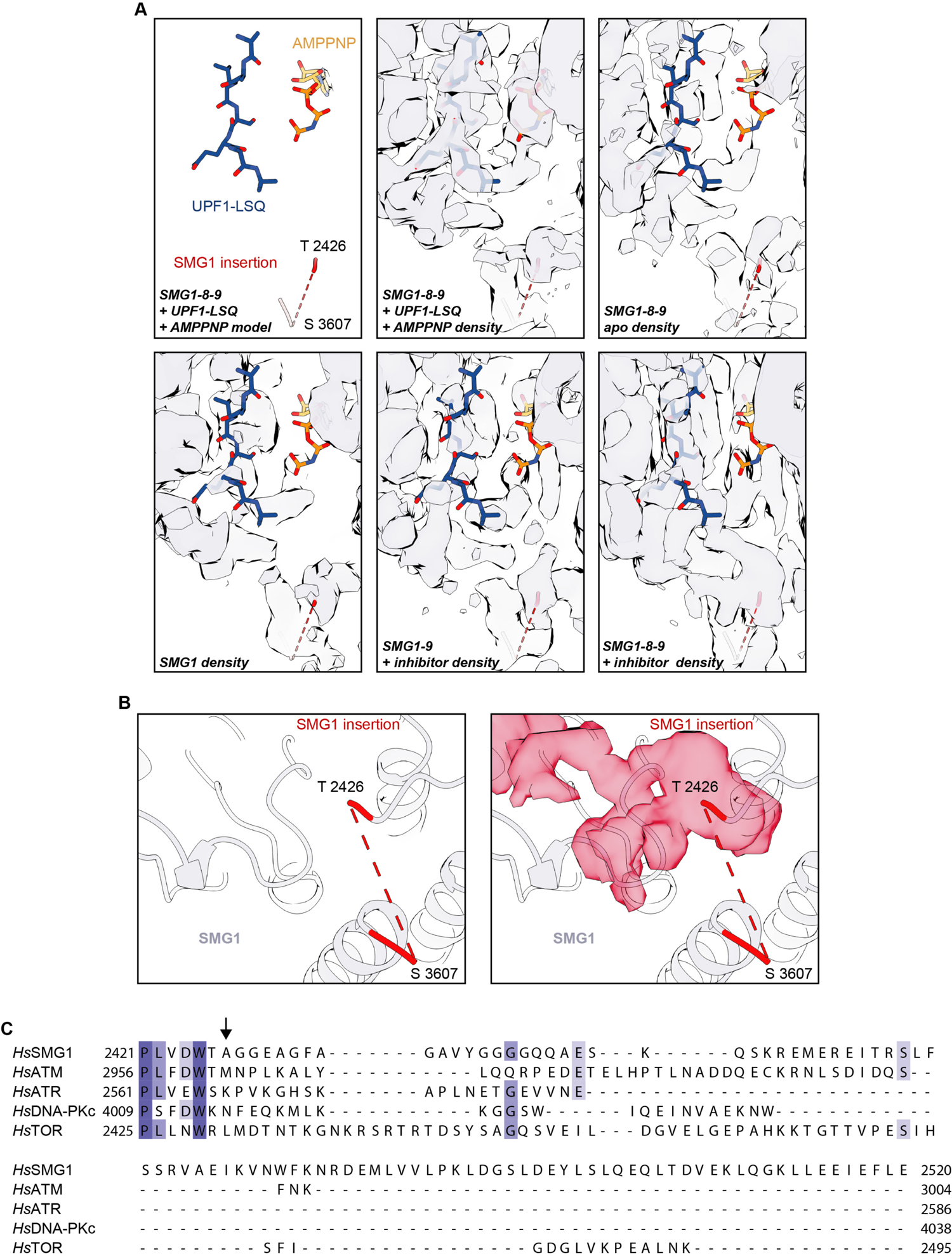
Details of the SMG1 insertion N-terminus. A) Model of SMG1 active site bound to UPF1-LSQ substrate and AMPPNP (PDB: 6z3r) shown superimposed with the corresponding EM density (EMD-11063) and the densities for apo SMG1-8-9 (EMD-10347), SMG1 (EMD-0836), SMG1**i**-bound SMG1-9 and SMG1**i**-bound SMG1-8-9. Note that the UPF1-LSQ model is partially covered by density protruding from the SMG1 insertion domain residue Thr 2426 only in the inhibitor-bound SMG1-8-9 complex. B) The isolated extra density (shown in red) observed in the active site of the SMG1**i**-bound SMG1-8-9 complex connects to the modeled N-terminus of the SMG1 insertion domain. The left panel highlights the last modeled N- and C-terminal residues of the insertion domain. The right panel shows the same view, superimposed with the isolated extra density. C) Multiple sequence alignment of the N-terminal 100 residues of the SMG1 insertion domain and the PRDs of the other human PIKK family members colored by identity. The beginning of the unmodelled part of the SMG1 insertion is indicated by a black arrow (compare B).

**Figure 3 - Supp. 2:**
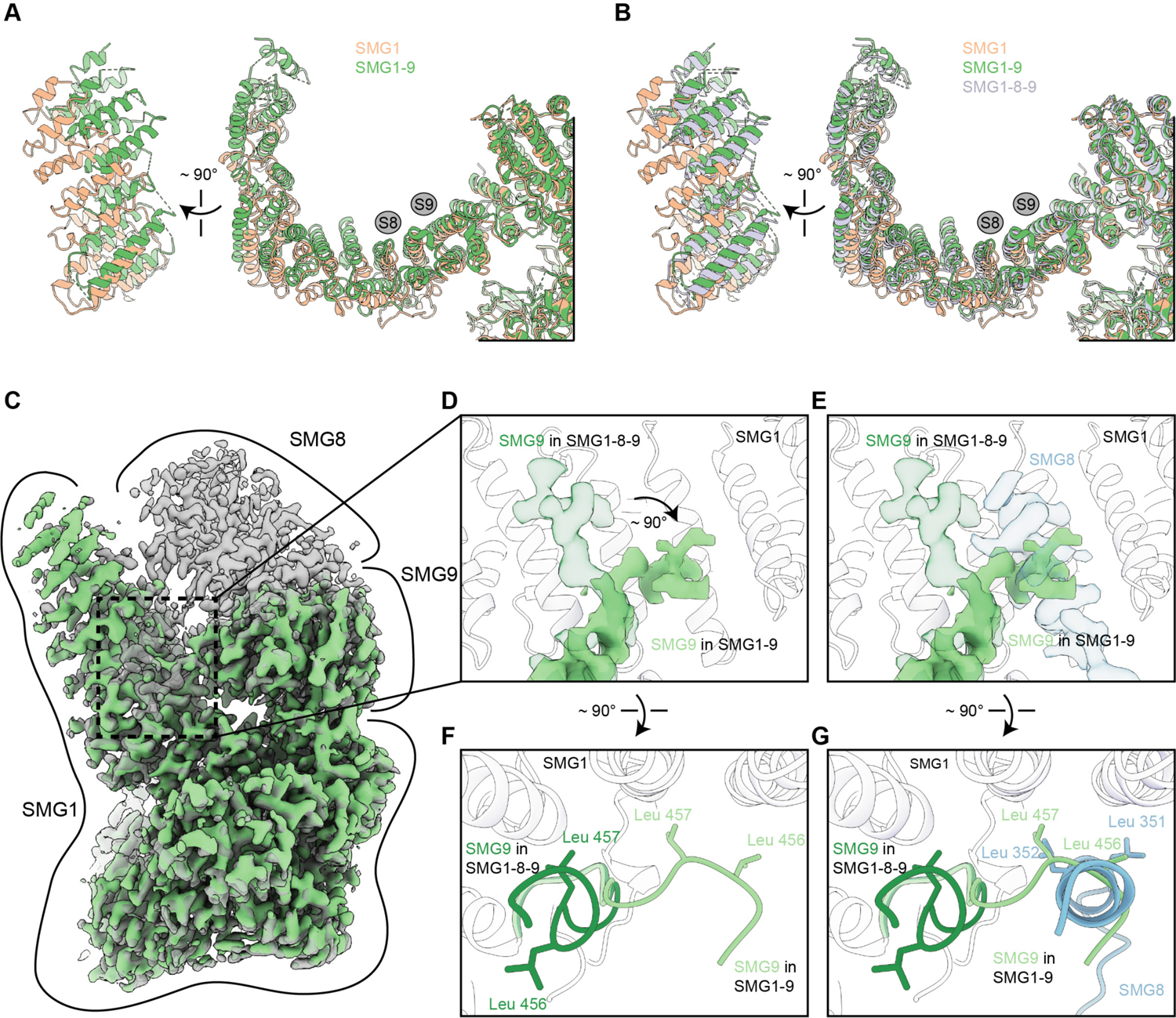
Details of the SMG1-9 complex. A) Overlay of SMG1 (PDB: 6l53) and SMG1-9 detailing movements of the N-terminal HEAT repeats. A front and a side view are shown and binding sites for SMG8 and SMG9 are indicated by grey circles. B) As in A), but including SMG1-8-9 (PDB: 6z3r). Models of SMG8 and SMG9 were occluded. C) Overlay of densities for SMG1-9 and SMG1-8-9 complexes, with SMG1-9 density in light green and SMG1-8-9 density in transparent grey. Approximate location of the single proteins within the densities are indicated. D) Close-up showing the rearrangement of a SMG9 segment in the SMG1-9 complex compared to the SMG1-8-9 complex. Density of the segment in SMG1-9 is shown in light green, superimposed with the SMG9 segment in SMG1-8-9 displayed in transparent dark green. The model for the interacting region of the SMG1 arch is shown as a transparent cartoon, and SMG8 is not shown. E) Same as D) but with density for a SMG1-interacting SMG8 segment in the SMG1-8-9 complex shown in transparent blue, that would clash with the SMG9 segment conformation observed in the SMG1-9 complex. F) Overlay of the model of the SMG9 segment in the SMG1-9 (light green) and the SMG1-8-9 (dark green) complex. Two Leu residues undergoing rearrangement between the two complexes are shown. Position of SMG1 is indicated, SMG8 is not shown. G) Same as in F) but shown with the SMG1-interacting SMG8 segment (blue). In the SMG1-9 complex, the highlighted pair of Leu residues in SMG9 substitutes for a pair of SMG8 Leu residues on the SMG1 arch.

**Figure 4 - Supp. 1:**
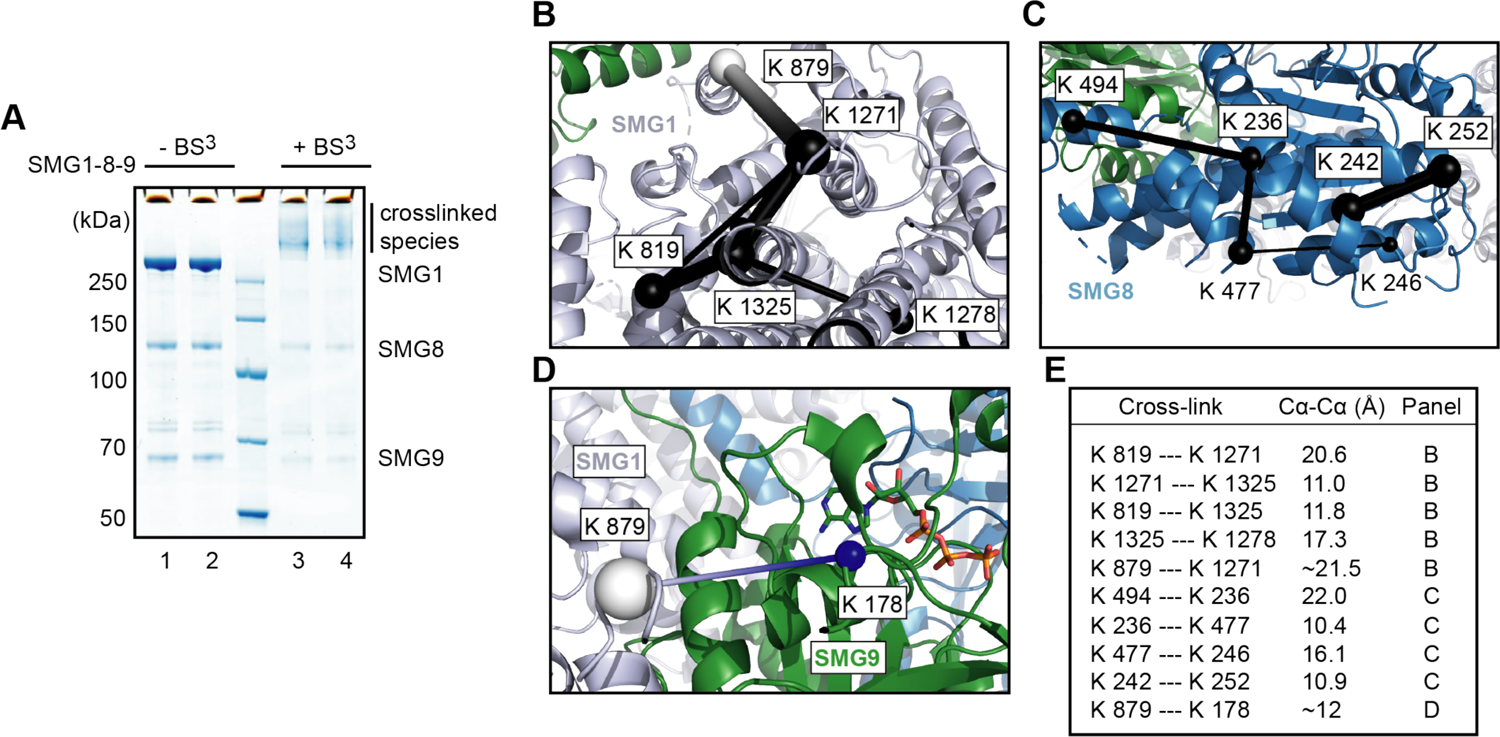
Cross-linking mass spectrometry of SMG1-8-9. A) Two samples of SMG1-8-9 (lanes 1 and 2) were incubated with BS^3^ (lanes 3 and 4) and analyzed using SDS-PAGE and Coomassie staining. B) Exemplary intra cross-links detected for SMG1 mapped on the model of the apo complex (PDB identifier: 6syt). Intra cross-links are shown in black with the thickness of the line indicating their score (thicker = higher score), and the respective residues indicated as spheres. A cross-link to an unmodeled region is shown in white (placed in the middle of the segment whose ends are the closest visible Cα atoms). C) Same as B), but for SMG8. D) Inter cross-link between SMG1 and SMG9. The Mg^2+^-ion complexed with SMG9 was omitted. E) Table listing measured distances for cross-links visualized in panels B), C) and D).

**Figure 4 - Supp. 2:**
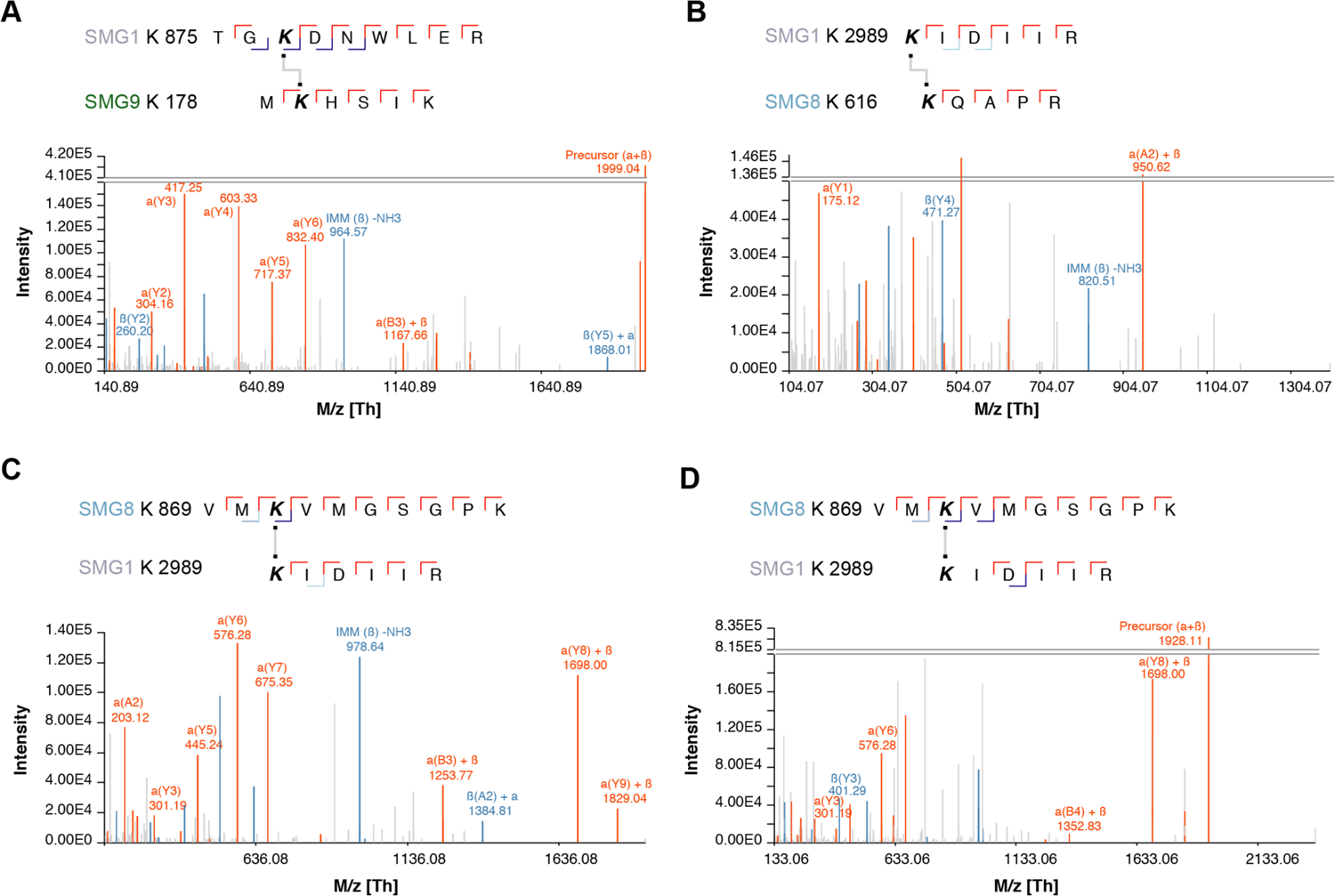
Selected spectra of detected SMG1-8-9 intra-links. A) to D) Cross-links are shown above each panel. All spectra showed good sequence coverage with full y-ion series, many b-ions and highly specific fragments.

**Figure 4 - Supp. 3:**
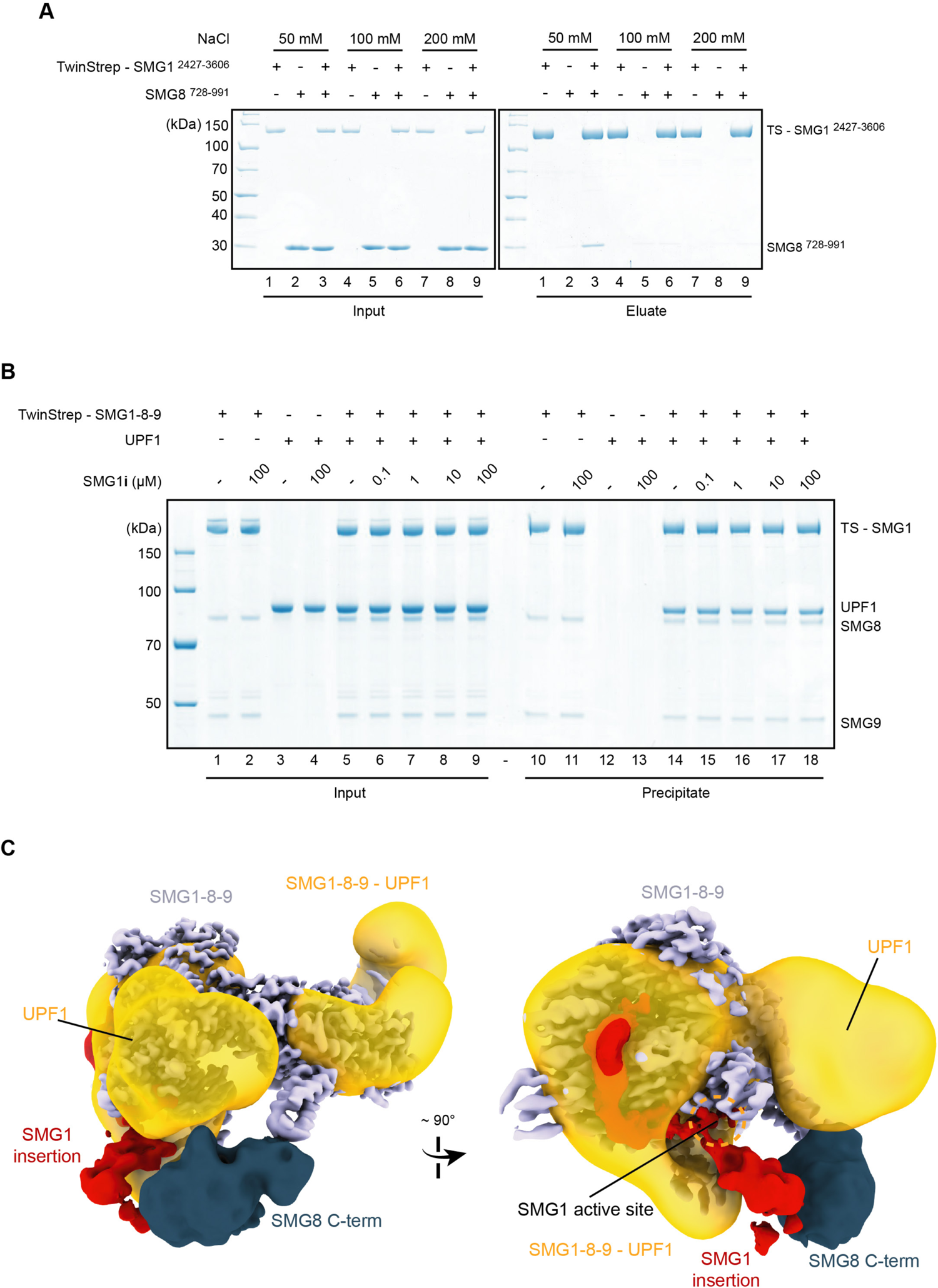
Further characterization of SMG1-8-9 - centered interactions. A) Coomassie-stained SDS-PAGE analysis of pull-down experiment showing that the interaction between SMG1 insertion domain (SMG1 ^2427-3606^) and SMG8 C-terminus (SMG8 ^728-991^) is dependent on low-salt conditions. B) Coomassie-stained SDS-PAGE analysis of pull-down experiment showing that SMG1-8-9 - SMG1**i** complex formation does not prevent interaction with UPF1. 0.12 μM of TS-SMG-1-8-9 and 0.4 μM of UPF1 were used throughout. C) The resolution-filtered, segmented cryo-EM density of SMG1**i**-bound autoinhibited SMG1-8-9 complex fitted in a negative-stain reconstruction of a cross-linked SMG1-8-9 - UPF1 complex (EMD-2664) shown in orientations similar to Fig. 4 A. The density within the negative-stain reconstruction assigned to UPF1 is indicated.

**Supplemental Table 1:** Cryo-EM data collection, refinement and validation statistics.

|  | SMG1 body<br>+ SMG1i | SMG1-8-9 +<br>SMG1i | SMG1-9 +<br>SMG1i | SMG1-8-9<br>+ AMPPNP | SMG1-9 +<br>AMPPNP |
| --- | --- | --- | --- | --- | --- |
| **Data availability** |  |  |  |  |  |
| EMDB | XXXX | XXXX/XXXX | XXXX | XXXX | XXXX |
| PDB | XXXX | XXXX/- | XXXX | XXXX | XXXX |
| **Data collection and processing** |  |  |  |  |  |
| Microscope | FEI Titan<br>Krios GII |  |  |  |  |
| Voltage (kV) | 300 |  |  |  |  |
| Camera | Gatan K3 |  |  |  |  |
| Energy filter | Gatan<br>Quantum-LS<br>(GIF) |  |  |  |  |
| Magnification | 105,000x |  |  |  |  |
| Pixel size (Å/pix) | 0.8512 |  |  |  |  |
| Target defocus range<br>(µm) | -0.5 to -2.5 |  |  |  |  |
| Electron exposure (e<sup>-</sup>/Å<sup>2</sup>) | 89.32 | 89.32 | 89.32 | 60.99 | 60.99 |
| Number of movies | 8,784 | 8,784 | 8,784 | 12,117 | 12,117 |
| Initially selected particle<br>candidates | 1,898,472 | 1,898,472 | 1,898,472 | 2,359,962 | 2,359,962 |
| Final number of particles | 228,291 | 228,291 | 85,216 | 403,079 | 168,050 |
| Resolution<sub>FSC independent</sub><br><sub>halfmaps</sub> (Å)<sup>a</sup> | 3.05 | 3.27 | 3.69 | 2.82 | 3.12 |
| **Refinement** |  |  |  |  |  |
| Initial model used | 6z3r | 6z3r | 6z3r | 6z3r | 6z3r |
| No. atoms | 10,396 | 18,973 | 16230 | 18,965 | 16342 |
| Residues | 1416 | 2630 | 2251 | 2629 | 2261 |
| Ligands |  |  |  |  |  |
| SMG1i | 1 | 1 | 1 | - | - |
| AMPPNP | - | - | - | 1 | 1 |
| ATP | 1 | 1 | 1 | 1 | 1 |
| IP6 | 1 | 1 | 1 | 1 | 1 |
| Mg | 1 | 1 | 1 | 1 | 1 |
| CC<sub>mask</sub>, CC<sub>box</sub>, CC<sub>peaks</sub>,<br>CC<sub>volume</sub><sup>b</sup> | 0.82, 0.79,<br>0.73, 0.79 | 0.82, 0.79,<br>0.71, 0.80 | 0.76, 0.76,<br>0.65, 0.73 | 0.82, 0.78,<br>0.75, 0.79 | 0.79, 0.73,<br>0.67, 0.76 |
| Resolution<sup>a</sup> FSC map vs. model<br>(0/0.143/0.5) (Å)<sup>b</sup> | 2.5/2.9/3.1 | 3.0/3.0/3.3 | 3.3/3.4/3.8 | 2.1/2.5/2.7 | 2.9/3.0/3.3 |
| *B* factors (Å<sup>2</sup>) |  |  |  |  |  |
| Protein | 21.99 | 80.93 | 71.54 | 20.68 | 44.18 |
| Ligands | 31.88 | 72.59 | 67.59 | 61.23 | 37.92 |
| RMS deviations |  |  |  |  |  |
| Bond lengths (Å) | 0.002 | 0.002 | 0.002 | 0.002 | 0.002 |
| Bond angles (°) | 0.543 | 0.529 | 0.527 | 0.631 | 0.562 |
| **Validation** |  |  |  |  |  |
| Ramachandran plot |  |  |  |  |  |
| Favored (%) | 97.28 | 97.42 | 97.27 | 96.38 | 96.97 |
| Allowed (%) | 2.72 | 2.58 | 2.73 | 3.17 | 2.99 |
| Disallowed (%) | 0.00 | 0.00 | 0.00 | 0.00 | 0.05 |
| MolProbity score | 1.33 | 1.34 | 1.41 | 1.38 | 1.45 |
| Clash score | 4.17 | 4.45 | 5.19 | 4.02 | 5.15 |
| Rotamer outliers (%) | 0.10 | 0.00 | 0.00 | 0.06 | 0.00 |

<sup>a</sup>according to the Fourier Shell Correlation (FSC) cut-off criterion of 0.143 defined in (Rosenthal and Henderson, 2003)
<sup>b</sup>according to the map-vs.-model Correlation Coefficient definitions in (Afonine et al., 2018a)

